# Neural divergence and convergence for attention to and detection of interoceptive and somatosensory stimuli

**DOI:** 10.1101/2020.06.04.134288

**Authors:** Aleksandra M. Herman, Clare E. Palmer, Ruben T. Azevedo, Manos Tsakiris

## Abstract

Body awareness is constructed by signals originating from within and outside the body. How do these apparently divergent signals converge? We developed a signal detection task to study the neural convergence and divergence of interoceptive and somatosensory signals. Participants focused on either cardiac or tactile events and reported their presence or absence. Beyond some evidence of divergence, we observed a robust overlap in the pattern of activation evoked across both conditions in frontal areas including the insular cortex, as well as parietal and occipital areas, and for both attention and detection of these signals. Psycho-physiological interaction analysis revealed that right insular cortex connectivity was modulated by the conscious detection of cardiac compared to somatosensory sensations, with greater connectivity to occipito-parietal regions when attending to cardiac signals. Our findings speak in favour of the inherent convergence of bodily-related signals and move beyond the apparent antagonism between exteroception and interoception.

## Introduction

Bodily self-consciousness depends on the perception and awareness of bodily signals. It is a multidimensional concept including identification with one’s body (i.e. body-ownership), self-location of body and body parts in space, and the first-person perspective (Blanke, 2012; Park & Blanke, 2019). Although we tend to take the ability to become aware of and identify with our body for granted, bodily self-consciousness can be easily malleable as it relies on the brain’s ability to integrate online information about the body originating from different sensory modalities (Aspell, Lenggenhager, & Blanke, 2012; Ehrsson, 2012; Park & Blanke, 2019; Sel, Azevedo, & Tsakiris, 2017; K. Suzuki, Garfinkel, Critchley, & Seth, 2013; Tsakiris, 2010; Tsakiris & Haggard, 2005). Importantly, at any given moment in time during wakefulness the brain integrates interoceptive (i.e. internal sensory information originating from visceral organs signalling the internal state of the body, for example information regarding cardiovascular, respiratory or gastrointestinal system), exteroceptive (i.e. sensory information about external, environmental features, events or stimuli, provided by touch, vision, or audition) and proprioceptive information (originating from receptors in muscles and ligaments signalling the position of body parts in space).

To give an example that illustrates the cross-talk between sensory modalities and their importance for bodily self-consciousness, consider the Rubber Hand Illusion (RHI) where synchronous exteroceptive visuo-tactile stimulation between a rubber hand and the participant’s hidden hand typically results in subjective feelings of ownership for the rubber hand (Botvinick & Cohen, 1998). An important behavioural outcome of the RHI is a change in proprioception, that is, in the felt location of the participant’s real hand. More recent studies have also shown how interoceptive signals contribute to the experience of body-ownership. Participants with lower interoceptive accuracy, as measured by the heartbeat counting task (Schandry, 1981), report a greater subjective experience of the illusion, compared to individuals with higher interoceptive accuracy (Tsakiris, Tajadura-Jiménez, & Costantini, 2011). Interoceptive inputs during the task also affect the illusion, for example, visual feedback of participant’s own heartbeats, increased self-identification with the virtual body (Aspell et al., 2013; K. Suzuki et al., 2013). Similarly, synchronous affective touch, an interoceptive modality of affective and social significance, increases the experience of the RHI (Crucianelli, Krahé, Jenkinson, & Fotopoulou, 2018). Therefore, higher interoceptive accuracy (i.e. better ability to feel internal bodily sensations) makes one less susceptible to embody foreign objects, while simultaneous visual feedback of one’s heartbeat or affective touch, helps to accept such objects as part of one’s body.

Therefore, given the importance of interoceptive, proprioceptive, and exteroceptive inputs for body-representation (Ponzo, Kirsch, Fotopoulou, & Jenkinson, 2018; Stone, Keizer, & Dijkerman, 2018; Tsakiris, 2010; Tsakiris et al., 2011), embodiment and self-conscious awareness (Arzy, Thut, Mohr, Michel, & Blanke, 2006; Lou et al., 2004), it is crucial to understand how such sensory information are processed in divergent or convergent ways in the brain and are brought to awareness.

Past neuroimaging research on the neural correlates of interoception has primarily assessed attention to cardiac activity (Avery et al., 2014; Caseras et al., 2013; Critchley, Wiens, Rotshtein, Ohman, & Dolan, 2004; Kuehn, Mueller, Lohmann, & Schuetz-Bosbach, 2016; Pollatos, Schandry, Auer, & Kaufmann, 2007; Simmons et al., 2013; Stern et al., 2017; Wiebking et al., 2010; Wiebking & Northoff, 2015; Zaki, Davis, & Ochsner, 2012), with a growing interest in respiratory-focused interoception (Farb, Segal, & Anderson, 2013; Wang et al., 2019) and sensations from the gut (Simmons et al., 2013). Typically, in these studies an interoceptive condition (sensing the internal state of the body; Craig, 2002) is contrasted against an exteroceptive condition (sampling the external world) using, for example, auditory (Caseras et al., 2013; Critchley et al., 2004; Kuehn et al., 2016; Pollatos et al., 2007; Wiebking et al., 2010; Wiebking & Northoff, 2015; Zaki et al., 2012) or visual stimuli (Avery et al., 2014; Simmons et al., 2013; Stern et al., 2017; Wang et al., 2019). Across these studies we observe very similar activation patterns for interoceptive vs control contrasts, pointing to increased activation of several cortical regions including the insular cortex, sensorimotor regions (postcentral gyrus, inferior parietal lobule, paracentral lobule, precentral gyrus, supplementary motor area) as well as occipital and temporal cortices, anterior cingulate, and lateral prefrontal regions during interoceptive condition. The insular cortex, particularly the right anterior insular cortex, is considered the main hub of the interoceptive network (A. D. (Bud) Craig, 2003, 2009a; Critchley et al., 2004). A small meta-analysis on cardioception revealed that attention to heartbeats relative to exteroceptive attention most consistently activates bilateral insula as well as premotor regions (Schulz, 2016).

However, the boundary between interoceptive and exteroceptive sensations becomes less clear when considering more proximal senses such as taste (chemosensing stimuli entering the body), touch (feel things close to us or in contact with us through skin, require close proximity to the body to be sensed) or proprioception (internally generated signals concerning the position of the body in space), as opposed to more distal senses such as vision and audition, which do not require such close proximity from the body. Specifically, touch gives us information about the way the skin surface of our body is embedded in and interacts with the environment and is an integral part of the existential experience of being a physical creature (O’Shaughnessy, 1989). Vision, on the other hand, informs us mainly about the surroundings and is especially important when it comes to actively exploring and navigating in the world. Thus, vision or hearing can be considered *distant senses* while touch can be considered a *proximal sense* (Klatzky & Lederman, 2011; Rodaway, 2002).

Regarding the question of bodily self-consciousness, somatosensory and proprioceptive signals are thought to be experientially self-specific (i.e. they concern one’s own body) in ways that vision and audition are not. Beyond the phenomenal experience, different types of tactile signals are transmitted through proprioceptive, exteroceptive and interoceptive pathways (Liljencrantz & Olausson, 2014; Olausson et al., 2008; Roudaut et al., 2012). Various receptors and afferent fibres are engaged in tactile stimuli detection and transmission (Roudaut et al., 2012). For example, Ruffini corpuscles located in dermis detect skin stretch and movement direction, while Pacinian corpuscules detect vibration. Vibrotactile stimulation elicits activation of primary and secondary somatosensory cortex as well as insula and thalamus (e.g., Briggs et al., 2004; Chakravarty, Rosa-Neto, Broadbent, Evans, & Collins, 2009; Chang et al., 2009; Golaszewski et al., 2006; Nelson, Staines, Graham, & McIlroy, 2004). Affective touch, which conveys emotionally-valent information through low mechanical threshold unmyelinated C fibres, also projects to the insula (Björnsdotter, Morrison, & Olausson, 2010; Liljencrantz & Olausson, 2014; Olausson et al., 2008, 2002). However, even though both somatosensation and interoception provide information about the body which might be important for bodily self-consciousness, there is a knowledge gap on the degree of overlap between tactile exteroception and visceral interoception. Therefore, considering a more proximal sense such as somatosensation alongside interoceptive processing might lead to novel insights regarding how these two sides of embodiment converge or diverge in the brain.

Indeed, a recent meta-analysis of 40 studies assessed the neural networks associated with perception of bodily sensations: those coming from inside the body (i.e. interoceptive) as well as externally to the body (e.g. rubber hand illusion, body ownership, self-location studies) (Salvato, Richter, Sedeño, Bottini, & Paulesu, 2019). A variety of interoceptive channels besides cardioception were investigated, including sensations such as thirst, air-hunger, attention to spontaneous bodily sensations, affective touch, and gastric balloon distension. Interestingly, processing of stimuli of the two domains converged primarily in the supramarginal gyrus, the right precentral, postcentral, and superior temporal gyri. Therefore, overlapping neural networks are engaged in interoceptive and exteroceptive body-related processing contributing to the creation of a multidimensional representation of the bodily self (Salvato et al., 2019). Yet, to our knowledge, a comprehensive study looking at a direct comparison between attention to and perception of interoceptive and somatosensory sensations is missing.

Noteworthy, so far neuroimaging studies investigating the neural correlates of interoceptive processing have primarily focused on aspects of *interoceptive attention*, that is the ability to direct attentional resources towards the source of internal body sensations (Khalsa et al., 2018). Our knowledge of neural processes engaged in *interoceptive detection*, defined as the ability to consciously detect the presence or absence of a stimulus (Khalsa et al., 2018), is limited despite the growing evidence of the importance of interoceptive accuracy as well as preconscious impact of afferent signals in behaviour and cognition (Critchley & Garfinkel, 2017; Garfinkel & Critchley, 2016; Quadt, Critchley, & Garfinkel, 2018). In exteroceptive domains, a meta-analysis (Meneguzzo, Tsakiris, Schioth, Stein, & Brooks, 2014) of neuroimaging studies comparing neural correlates of supra- vs subliminal presentation of the same modality (visual, auditory, or tactile) revealed that conscious detection of the exteroceptive stimuli was associated with greater activity in left anterior cingulate cortex and mid-caudal anterior cingulate cortex. Subliminal presentation (i.e. non-conscious perception), on the other hand, evoked consistently greater activations in the right fusiform gyrus/middle occipital gyrus, right caudal anterior cingulate cortex and right insula. Therefore, anterior cingulate cortex was most consistently activated in response to both subliminal and supraliminal stimuli presentation, presumably playing a role in integration of conscious and non-conscious processing (Meneguzzo et al., 2014). In the interoceptive domain, Critchley and colleagues (Critchley et al., 2004) utilised a heartbeat discrimination task (Whitehead, Drescher, Heiman, & Blackwell, 1977), whereby participants are asked to judge whether a series of tones is presented in sync with one’s heartbeats (presented at cardiac systole) or delayed (presented at cardiac diastole). This task involves correct detection of internal signals (heartbeats) and an ability to differentiate them from external stimuli (tones). However, the exteroceptive control task is different: participants need to judge whether all tones in a series are the same or whether one is different (odd-one-out). Thus, these tasks likely involve different processes. Most commonly used heartbeat counting task (Schandry, 1981), on the other hand, requires participants to silently count their own heartbeats in predefined periods. Performance in this task, however, can be affected by various factors, including knowledge of one’s heart rate or counting seconds instead of heartbeats and its validity has recently been criticised (e.g. Ring & Brener, 2018; Zamariola, Maurage, Luminet, & Corneille, 2018; also see Ainley, Tsakiris, Pollatos, Schulz, & Herbert, 2020 for further discussion). Moreover, using these tasks, we cannot differentiate between neural activation when attending to vs conscious detection of a stimulus. Investigating the neural correlates of conscious detection of heartbeats requires the use of a task that allows to reliably dissociate between instances of detected and attended but not detected heartbeats.

Given the recent interest in neurocognitive models of bodily self-consciousness (Blanke, 2012; A. D. (Bud) Craig, 2009b; Tsakiris, 2017) and the existing literature on how somatosensation and interoception are cortically represented (Salvato et al., 2019), we set out to investigate the potentially divergent and convergent ways in which attention to and detection of somatosensory and interoceptive signals are processed. Thus, the aim of the current study was to identify and compare the neural correlates of directed attention as well as conscious and non-conscious perception of heartbeats and tactile (somatosensory) stimuli. To do this we employed an MRI compatible ECG system in order to accurately align heartbeats to the fMRI signal and designed a novel Heartbeat/Somatosensory Detection task in order to dissociate between felt and not felt stimuli during an fMRI scan. We tested three hypotheses: (1) *attention* to interoceptive and somatosensory stimuli would yield overlapping but dissociable activation patterns across the brain (e.g. insula cortex, somatomotor cortex, and thalamus); (2) conscious *detection* of interoceptive and somatosensory sensations would yield overlapping, but dissociable activation patterns across the brain; and (3) as the central hub of the interoceptive network (A. D. (Bud) Craig, 2003; Critchley et al., 2004), but also a crucial part of the cognitive-control and salience processing network (Jiang, Beck, Heller, & Egner, 2015; Uddin, 2015; Wang et al., 2019), functional connectivity with the right insular cortex would be modulated by conscious detection of stimuli across interoceptive and somatosensory conditions. Thus, our study goes beyond past investigations as it addresses the independence and overlap of directed attention to interoceptive and somatosensory cues, as well as contrasting the neural correlates of conscious and non-conscious processing of these stimuli.

## Methods

We report how we determined our sample size, all data exclusions, all inclusion/exclusion criteria, whether inclusion/exclusion criteria were established prior to data analysis, all manipulations, and all measures in the study.

### Participants

38 participants in total (aged 19-52, 26.4±6.94; 16 males) were recruited for the study and completed a first behavioural screening session. Participants were selected for the MRI scan based on their ability to subjectively feel their heartbeats in the Heartbeat Detection Task (see below). Participants completed a practise version, with 2 blocks of 20 trials each, of the experimental task to be carried out in the scanner in the behavioural screening session. Only those who felt their heartbeat on 40-80% of trials were invited to participate in the MRI session. This screening procedure ensured that participants scanned would have a distribution of both detected and undetected heartbeats. Thirty participants (aged 19-52, 26.83±6.82; 12 males) passed the screening and completed the MRI scan on a different day. The sample size was estimated based on previous research employing cardioceptive tasks in the fMRI environment (Farb et al., 2013; Stern et al., 2017; Wiebking et al., 2011), but not formal power calculation was performed. All participants provided written informed consent in line with the Local Ethics Committee Regulations and MRI Safety Procedures. At the time of testing, none of the participants were taking any medication for a neurological or psychological disorder or showed any MRI contradictions. Participant were asked to refrain from taking any caffeine three hours before the MRI scan.

For two participants the automatic detection algorithm was unable to detect any R peaks after pre-processing the ECG data from these blocks due to low signal-to-noise-ratio during the recording of the ECG. As those two individuals were removed from the analysis entirely due to poor ECG quality during MRI session, the final sample consisted of 28 participants. 25 of them had complete datasets (8 blocks), while the remaining three had seven blocks only, due to poor ECG quality or excessive motion (see below for details).

### Experimental Design

#### Heartbeat and Somatosensory Detection Task

Participants completed a novel Heartbeat and Vibrotactile Detection Task in the MRI scanner. The task was programmed in Cogent toolbox (Wellcome Dept., London, UK) for MATLAB 2015b (Mathworks Inc.). The experimental task was divided into two block types: heartbeat detection and somatosensory detection. At the beginning of each block, participants were instructed to either focus on their heart beating or detect a faint vibration presented on their left hand. The vibrotactile stimulator was secured to the skin above the first dorsal interosseous. The somatosensory stimuli, with a sinusoidal wave form of adjustable amplitude and of 150ms in duration, were delivered using MRI-compatible pneumatic vibrotactile device (dual channel vibrotactile transducer with MRI compatible tactile transducer system). On each trial, participants were presented with a black fixation cross for a pseudorandomised inter-trial interval (ITI) ranged from 4 to 8 seconds in 10 steps. Each trial consisted of three epochs, whereby the fixation cross changed colour from red to green to blue (750ms each) followed by a response screen (see Fig 1 for a schematic). While the response screen was presented, participants were instructed to press the button (or buttons) corresponding to the colour of the cross during which they felt a target sensation (heartbeat or somatosensory). It was emphasised that they should take a conservative approach and provide a button press when they actually felt the sensation, i.e. not to guess on any instance, but also that they could press multiple buttons depending on when they felt a stimulus (i.e. during which colourful cross). If they did not feel anything, they pressed the “NO” button. This ensured a button was pressed following every trial. Another response screen followed, during which participants rated their confidence in the response on a scale of 1-4. If participants indicated that they felt a stimulus, the response screen asked how confident participants were that they had felt a stimulus; however, if participants indicated that they did not feel a stimulus, the response screen asked how confident participants were that they had not felt a stimulus. Both response screens were presented for a fixed time of 2500ms. This was to ensure that trials remained as consistent as possible across conditions.

**Figure 1.**
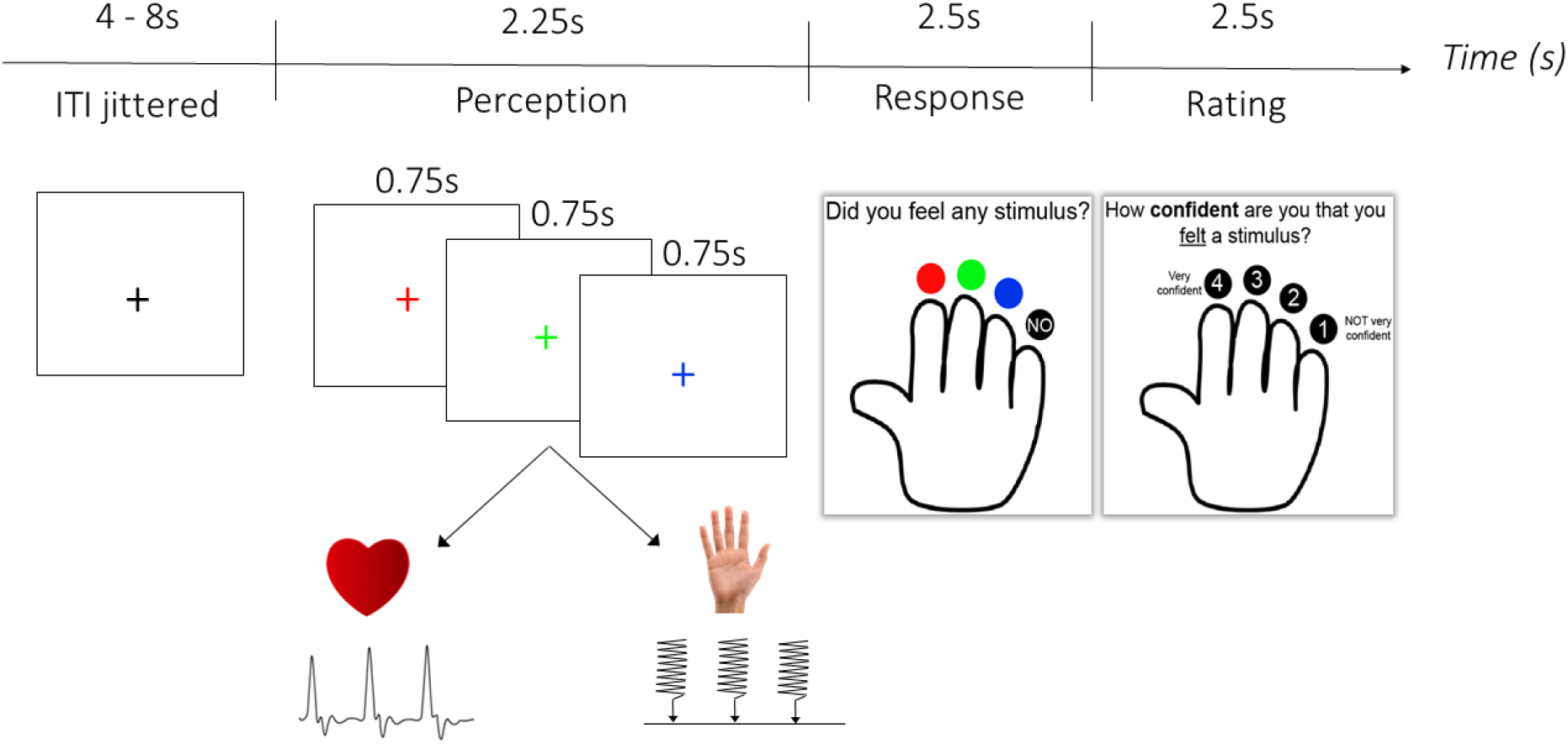
A schematic of a single trial in the Heartbeat and Somatosensory Detection Task. Participants were first instructed to focus either on their heart beating or faint vibrations applied to their left hand. Following a variable inter-trial interval (ITI), fixation cross changed colour three times (stimuli perception phase). Next, participants had to indicate with a corresponding button press (or multiple button presses), when (i.e. during presentation of which of the fixation crosses) they felt a stimulus/stimuli. A button press was also required if participant did not feel any stimuli within the perception phase. Finally, they rated their confidence.

Importantly, as participants’ hearts were beating continuously throughout the experiment, to maintain the same sensory stimulation as much as possible between conditions, somatosensory stimuli were also presented on the left hand continuously throughout all blocks. The inter-stimulus interval (ISI) was set to match the participants’ heart rate as closely as possible and some pseudorandomised variation was added to the ISI between 0 and 90ms to ensure this did not become too predictable and mirror typical heart-rate variability. Importantly, the ISI was based on heart-rate, but no time-locking to the ECG signal was done (i.e. vibrotactile stimuli could occur at any point in the cardiac cycle). To maximise the match between the conditions, the intensity of stimuli presentation was set to just below the individual somatosensory perception threshold (see below) with some occasional fluctuations above the threshold. Participants completed 8 blocks in total (4 of heartbeat detection and 4 of somatosensory detection) with 20 trials per block (60 epochs). The block type was alternated with the order counterbalanced across participants.

#### Somatosensory Thresholding Procedure

Before starting the main task, participants completed a thresholding task to calibrate the intensity of the somatosensory stimulation. The task was exactly the same as the main task (to allow sufficient practise on the task), however, only a single somatosensory stimulus was presented on each trial and participants reported when they felt it.

The intensity of the tactile stimulus was controlled using the volume of a sound system attached to the tactile device. The amplitude of the sound waves was converted into an air puff of a given intensity. At the beginning of the experiment this volume was always set to its maximum. An initial tuning determined a rough estimate of the intensity in which the participant could just detect the tactile stimulus. This estimate was used as a prior in the Bayesian thresholding procedure employed using the QUEST toolbox in MATLAB (Pelli, 1987; Watson & Pelli, 1983). On each trial the probability density function (PDF) of the intensity was updated using the response on that trial. A new test intensity was then suggested as the best quantile of the posterior PDF. At the end of 40 trials, the final intensity estimate used was the mean of the posterior PDF. This follows the procedure outlined in Pelli (1987). The experimenter then analysed a plot of the changing intensity over trials to determine that the procedure converged on a stable estimate that did not continue to increase or decrease by multiple steps in the final 10 trials. If this was not the case the procedure was repeated until the experimenter was satisfied the procedure had converged on a stable estimate. The procedure was set to determine a threshold at which the participant detected the stimulus on 60% of trials. The threshold is expressed as a proportion of the maximum volume of the sound system. Using this procedure, the intensity of the tactile stimulus was standardised across individuals.

Throughout somatosensory detection blocks in the MRI scanner, the intensity of the somatosensory stimulus was monitored and modulated online using a staircase procedure to ensure that participants’ somatosensory detection was roughly at 50% in each block. Specifically, if the tactile proportion became greater than ~80% or less than ~40% in a tactile block then the intensity of the tactile stimulus was adjusted by 0.5. This was to try to ensure that the perceived intensity remained similar throughout the task even when the stimulus became predictable and was therefore more difficult to perceive. Any adjustments were made for pairs of blocks such that there was always a matching cardiac detection block with the same tactile intensity. No changes were made to the intensity of the tactile stimulus following a cardiac detection block.

#### Heartbeat Counting Task

During the behavioural screening session participants completed the heartbeat counting task (Schandry, 1981). Participants were asked to count how many heartbeats they could feel in a given period (25s, 30s, 35s, 40s, 45s, ad 50s, in a randomised order). The instructions were as follows: “Please sit back and relax and try to feel your heart beating in your chest. When you hear the start signal (auditory beep) please start counting your heartbeats and stop when you hear the stop signal (auditory beep). You can have your eyes open or closed during the task.” After inputting the number of heartbeats counted on each trial, participants rated how confident they were in their answer on a scale of 0-100. Participants completed six trials.

The dependent variable of the heartbeat counting task is the interoceptive accuracy (IAcc) score, which serves as an objective measure of how well an individual can feel their heart beating (Schandry, 1981). IAcc is calculated by determining the proportion of counted heartbeats over actual heartbeats on each trial and then averaging this over trials and deducting from 1 using the following formula: 1-[(∑N(counted beats / actual beats))/N], where ‘N’ equals number of trials.

### Data collection

All MRI data was collected in a Siemens Magnetom TrioTim syngo MR B17 3-Tesla scanner (Siemens AG, Munich, Germany) at the CUBIC imaging centre at Royal Holloway, University of London.

First, structural volumes were obtained using the high-resolution three-dimensional magnetization rapid acquisition gradient echo sequence (TR = 1.9 s, TE = 3.03 ms, TI = 1.1 s, FA = 11°, 144 sagittal slices per slab, 1 × 1 × 1 mm, FoV = 256 mm, GRAPPA acceleration factor = 2). Next, whole-brain multiband gradient echo echo-planar imaging (EPI) sensitive to blood oxygenation–level dependent signal was used to collect fMRI data (multiband acceleration factor = 2, TR = 1100 ms, TE = 30 ms, FA = 76°, 32 slices, FoV = 192 mm, voxel size = 3 × 3 × 3 mm, 5:03 min/block). After 4 blocks of the task, whilst participants rested, a fieldmap was acquired using the same resolution and slice locations as multiband images, to allow for offline correction of field inhomogeneities (TR = 525 ms, TE = 5.19/7.65 ms, FA = 60°, 1:10 min).

Throughout the MRI scan, we collected electrocardiogram (ECG) data using MRI compatible ECG electrodes and leads (BIOPAC). These were configured in a tight right-angled triangle on the left side of the chest. The skin was scrubbed using an abrasive cloth and prepped using Nuprep Skin Prep Gel (D.O. WEAVER and COMPANY) before the electrodes were attached. The ECG signal was recorded with a Powerlab 8/35 box (Bio Amp 132) and LabChart 8 software (www.adinstruments.com).

### Data Analysis

#### ECG data

Due to the artefacts from the EPI sequence, the ECG data required a large amount of preprocessing to extract timing of each R peak during the task. This was completed using in-built functions within Acqknowledge software (BIOPAC). The ECG data was filtered sequentially at 50Hz and 14.54Hz (EPI scanner frequency) using a comb band stop filter. A window of 600-900ms (depending on heart rate) was selected around heartbeats prior to the start of the EPI sequence. These epochs were averaged to create a QRS template. A normalised cross-correlation then correlated this template with the whole ECG timeseries in an overlapping sliding window. Peaks greater than 0.5 correlation were detected and labelled as QRS complexes then superimposed onto the filtered ECG trace. Each timeseries was then visually inspected and any missed or incorrectly labelled QRS peaks were manually edited.

The ECG quality was insufficiently good for two participants to reliably establish timing of the R-peaks; therefore, data from these two individuals was excluded from the analysis entirely. For an additional two participants, the ECG quality was poor for one of the Heart blocks; these blocks were also removed from the further analysis.

#### Behavioural Data Analysis

The main dependent variable for the experimental task in the scanner was the participants’ response of feeling or not feeling the stimuli. For each trial, each coloured cross was treated as a separate epoch creating 60 epochs per block (20 trials). As per signal detection theory, each epoch was categorised as either a Hit, Miss, False Alarm or Correct Rejection depending on whether the participant indicated that they felt or did not feel a sensation during each epoch and whether the heartbeat or somatosensory stimulus was present or absent. To quantify the performance, we calculated an accuracy score [Accuracy = (N_Hits_ + N_Correct rejections_)/N_epochs_] for each block and condition. For completeness, we also calculated d’ as a signal detection theory index of individual sensitivity to heartbeats and somatosensory stimuli. D’ was calculated taking all trials into account for Cardiac and Somatosensory Focus conditions separately. The performance on the task was analysed using a 2 (Cardiac vs Somatosensory condition) by 4 (blocks) repeated measures analysis of variance (rmANOVA) or paired-samples *t*-test, as appropriate, conducted in R implemented in R Studio (R Studio Team, 2016).

#### MRI Data

FMRI data pre-processing and analyses were carried out using FEAT (FMRI Expert Analysis Tool) Version 6.00, part of FSL (FMRIB’s Software Library; Jenkinson, Beckmann, Behrens, Woolrich, & Smith, 2012).

##### PRE-PROCESSING

Pre-processing steps included skull stripping of structural images with Brain Extraction Tool (BET; Smith, 2002), removal of the first four functional volumes to allow for signal equilibration, head movement correction by volume-realignment to the middle volume using MCFLIRT (Jenkinson, Bannister, Brady, & Smith, 2002), global 4D mean intensity normalization, spatial smoothing using a Gaussian kernel of FWHM 6mm, grand-mean intensity normalisation, high pass temporal filtering (Gaussian-weighted least-squares straight line fitting, with sigma=50.0s) and fieldmap based distortion correction. Participants’ motion was minimal and did not exceed 3 mm (1 voxel) with the exception of a single Heart Focus block for one of the participants where movement spikes exceeded this threshold. This run was, therefore, excluded from further fMRI analysis. Registration to high resolution structural images was carried out using FLIRT (Jenkinson et al., 2002; Jenkinson & Smith, 2001). Registration from high resolution structural to MNI152 standard space was then further refined using FNIRT nonlinear registration (Andersson, Jenkinson, & Smith, 2010).

##### UNIVARIATE ANALYSIS

Time-series statistical analysis was carried out using FILM with local autocorrelation correction (Woolrich, Ripley, Brady, & Smith, 2001). In the first-level modelling, customized waveforms (for each participant, run and event type) representing each event type onset and the duration of stimulus presentation were convolved with a double-gamma hemodynamic response function and a high pass filter was applied to remove low-frequency artefacts. Two separate analyses were performed. To investigate the neural correlates underlying attention to the heart and somatosensory stimuli, we modelled general attention to heartbeats/somatosensory stimuli, taking into account the whole duration of Cardiac/Somatosensory perception across epochs (duration=2.25s) with the onset at the first (red) fixation cross. To investigate the neural correlates of conscious and non-conscious detection of these sensations, we separated the individual epochs as independent events (duration=0.75s), and categorised them as either a Hit, Miss, False Alarm or Correct Rejection, to match the behavioural analysis. For the detection analysis, the events were modelled at the onsets of each epoch (each colourful fixation cross). The button press onsets as well as response screen and confidence screen were additionally included as regressors of no interest.

Next, we estimated each participant’s mean neural response during Cardiac/Somatosensory Focus (focus analysis) or Hits and Misses for Cardiac and Somatosensory conditions separately (conscious detection analysis). To this end, for each first-level FEAT output, the four blocks for respective condition were combined for each participant using a second-level fixed effects GLM to create averaged maps.

To identify brain regions recruited more in response to Cardiac relative to Somatosensory condition, a third-level whole brain voxel-wise GLM was conducted across all participants for each of the (second-level) contrasts of interest. This between-subject analysis was carried out using the FMRIB Local Analysis of Mixed Effects (FLAME; Woolrich, Behrens, Beckmann, Jenkinson, & Smith, 2004). Z (Gaussianised T/F) statistic images were thresholded non-parametrically using clusters determined by Z > 3.1 and a (corrected) cluster significance threshold of *p* = 0.05 across the entire brain (Worsley, 2001).

Overall, there were three contrasts of interest: (1) the main effect of focus condition (Cardiac Focus vs Somatosensory Focus), (2) the main effect correct signal detection (Hits vs Misses), and (3) the interaction effect (Cardiac Hits – Cardiac Misses vs Somatosensory Hits – Somatosensory Misses).

For completeness, we also conducted additional set of analyses, whereby as opposed to modelling the whole epochs, we modelled the onsets of the heartbeats and vibrotactile stimuli. The details of that analysis and results is reported in Supplementary Materials.

In all reported analysis, the Harvard-Oxford cortical and subcortical probabilistic atlases (Desikan et al., 2006; Frazier et al., 2005; Makris et al., 2006) were used to identify each region revealed.

##### CONJUNCTION ANALYSIS

To identify regions that show common activity in Cardiac and Somatosensory conditions, we conducted a formal conjunction analysis (Nichols, Brett, Andersson, Wager, & Poline, 2005) using FSL easythresh_conj function (FMRIB, Oxford, UK, Part of FSL - FMRIB’s Software Library, *p* < 0.05).

##### PSYCHO-PHYSIOLOGICAL INTERACTION ANALYSIS

To look at task-specific changes in the relationship between activity in an identified seed region and other areas of the brain (O’Reilly, Woolrich, Behrens, Smith, & Johansen-Berg, 2012), we conducted a context-dependent psychophysiological interaction analysis (gPPI; McLaren, Ries, Xu, & Johnson, 2012).

The seed region was defined using the cluster from the conjunction analysis, limiting it to the area which encompassed the right Insular cortex. The seed region of interest (ROI) mask from the conjunction analysis was first transformed to each individual participant’s functional native space, using inverse warping. Next, the average time courses of the ROI were extracted from motion-corrected, high-pass filtered image data (same pre-processing steps as outlined above) for each participant using fslmeants. The gPPI analysis was conducted using FSL’s FEAT. The task variables were convolved with a double-gamma hemodynamic response function, and temporal derivatives for the task variables were included in the model. The element-by-element products of the Insula ROI timeseries and the convolved task regressor (embodying the contrast of Hits and Misses) were added to the model along with the raw ROI timeseries together with the remaining task variables as in the main univariate analysis. A whole-brain contrast image for the gPPI was computed from this model and submitted for second- and third level group analyses described above. The gPPI was tested as a contrast between the two interaction regressor coefficients (i.e., Cardiac Hits vs Misses × Insula ROI – Somatosensory Hits vs Misses × Insula ROI) (McLaren et al., 2012; O’Reilly et al., 2012). Additionally, to understand the relationship between insula connectivity and task performance better, we performed the PPI analysis for Cardiac and Somatosensory conditions separately. We report this analysis in the Supplementary Materials.

No part of the study analyses was pre-registered prior to the research being conducted.

## Results

### Behavioural Results

Since one block of the Heart Focus condition was missing for two individuals, the sample in all behavioural analyses consisted of 26 individuals. First, as a means of general comparison of both conditions, we compared the percentage of epochs where the signal of interest (i.e., heartbeat or somatosensory stimulation) was present during the scanning session (Fig. 2A). RmANOVA revealed the main effect of condition [*F*(1, 25) = 24.61, *p* < .001, η^2^ = 0.051], with on average more somatosensory stimuli than heartbeats present (87.23±12.05 and 82.05±10.65, respectively). There was also a significant main effect of block [*F*(3, 75) = 3.79, *p* = .014, η^2^ = 0.005], as well as a condition by block interaction [*F*(3, 75) = 2.87, *p* = .042, η^2^ = 0.005], driven by a gradual decrease in heartbeats present across the Heart Focus blocks, due to a trend-level decrease in heart rate over time [*F*(3,75) = 2.32, *p* = .082, η^2^ = 0.007; Fig. 2B]. The occurrence of somatosensory stimulation, on the other hand, was relatively constant throughout the task. This is because for some participants/blocks we were not able to get a readable ECG signal during blocks due to interference from the MRI scanner, therefore could only estimate heart rate at offline post-processing the data. For those who did have a clear ECG signal, despite the scanner interference, we estimated heart rate in between blocks and adjusted the tactile ISI accordingly, but this was not possible for all participants/bocks. Thus, the rate of somatosensory stimulation did not always account for the (slight) decreases in HR over time.

**Figure 2.**
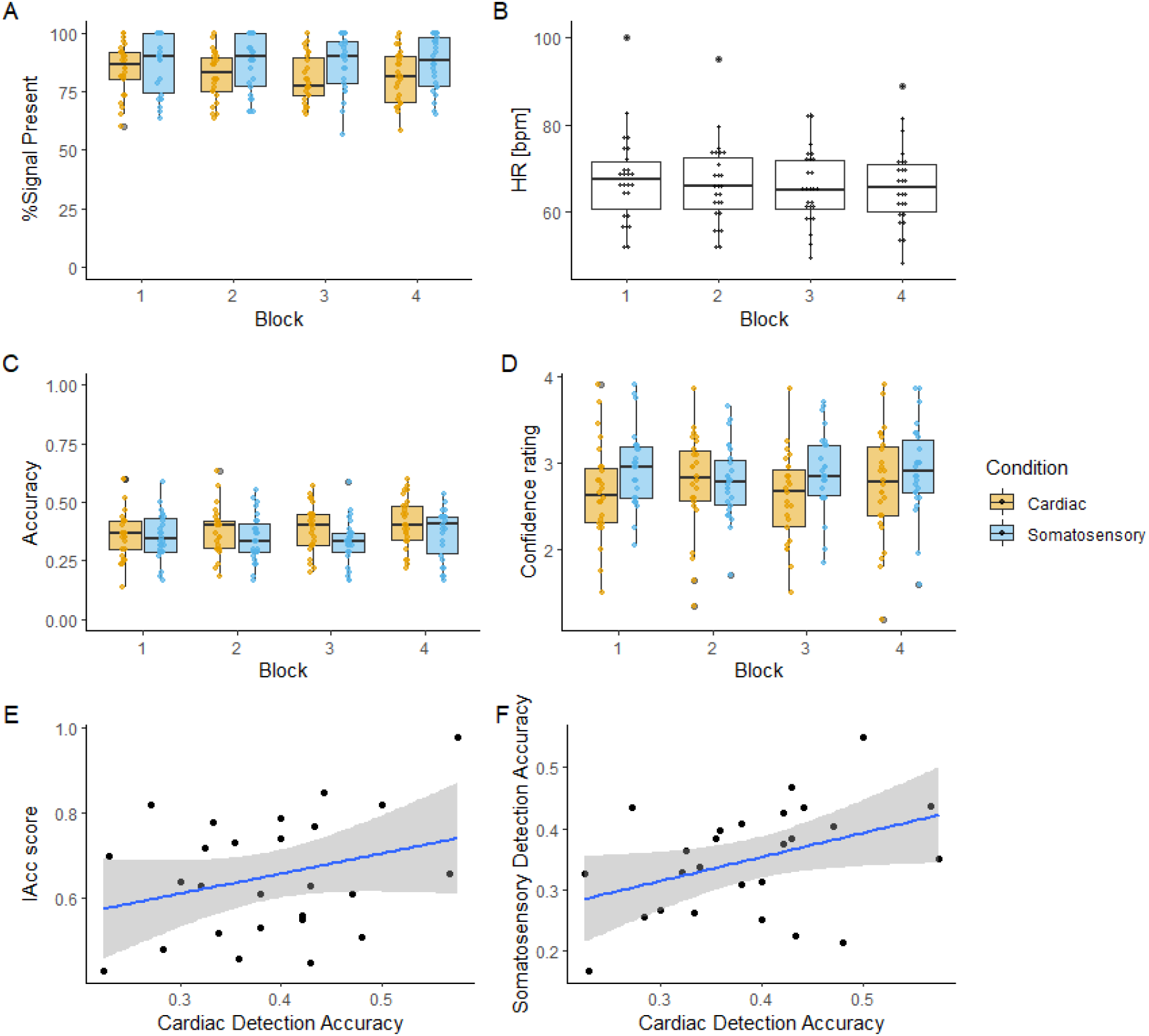
*Performance on the behavioural detection task during the scanning session. A. Percentage of epochs in which heartbeat or somatosensory stimuli were present. B. Average heart rate (HR) per Cardiac Condition block. C. Accuracy (proportion of Hits with Corrects Rejections) per block and condition. D. Mean confidence per block of the task conditions. E. Scatterplot presenting the relationship between the interoceptive accuracy (IAcc) score on the Heartbeat Counting Task and the accuracy on the Heartbeat detection Task [r(25) = 0.30,* p *= .133]. F. The relationship between accuracy on the Somatosensory and Heartbeat detection task [r(25) = 0.40,* p *= .043]. Shaded area in the scatterplots represents 95% CI*.

Secondly, we compared the accuracy on the task (the proportion of Hits + Correct Rejections). There was no significant main effect of condition [*F*(1, 25) = 3.99, *p* = .057, η^2^ = 0.034; Fig. 2C] although the effect was approaching significance with higher accuracy for the Heart vs Somatosensory Condition (0.39±0.09 vs 0.35±0.09, respectively). There was no main effect of block [*F*(3, 75) = 2.29, *p* = .085, η^2^ = 0.012] nor an interaction [*F*(3, 75) = 0.85, *p* = .471, η^2^ = 0.004]. For completeness, in Supplementary Materials we also present the proportion of hits and misses per condition and block as well as per epoch. We also calculated d’ as the signal detection theory index of sensitivity for all blocks collapsed together. As some participants did not have any false alarms we, therefore, calculated the d’ according to Hautus (1995) by adding 0.5 to each cell of the contingency table. The paired samples *t*-test revealed no significant differences in d’ between the focus conditions, *t*(27) = 0.10, *p* = .918, [−0.20, 0.22]. Finally, we used criterion as a signal detection theory index of a tendency to report that the signal was present. A larger value of the criterion in one condition would imply that stronger evidence for that condition is required before saying that the signal is present. The paired samples *t*-test, however, revealed no significant differences in criterions between the focus conditions, *t*(27) = 1.4, *p* = .173, [−0.26, 0.05], indicating that participants used comparable criteria to report that they feel a heartbeat and a somatosensory stimulus.

Additionally, we compared confidence ratings on the task (Fig. 2D). There was a main effect of condition [*F*(1, 25) = 7.83, *p* = .010, η^2^ = 0.032], with higher confidence for the Somatosensory (2.88±0.46) than the Cardiac (2.69±0.56) condition, no main effect of block [*F*(3, 75) = 1.02, *p* = .387, η^2^ = 0.003], but the interaction was significant [*F*(3, 75) = 3.76, *p* = .014, η^2^ = 0.011], suggesting that the confidence fluctuated differently across blocks for the Cardiac and Somatosensory Conditions.

Finally, to compare in-the-scanner task performance with the accuracy in the more-established Heartbeat Counting Task, which was carried out during the practise behavioural session outside of the scanner, we computed Pearson’s correlation coefficient between Accuracy in the Heartbeat Detection Task and IAcc score (Fig. 2E). We found a positive but not-significant relationship between the two measures, *r*(25) = 0.30, *p* = .133, suggesting that participants who performed well in the Heartbeat Detection Task did not necessarily have high accuracy in the Heartbeat Counting Task. There were also no significant correlations between IAcc and accuracy in the somatosensory detection condition of the in-the-scanner detection task, *r*(25) = 0.12, *p* = .575, but performance in the heart detection condition did correlate with performance in the somatosensory detection condition, *r*(25) = 0.40, *p* = .043 (Fig. 2F). Important to note that individuals for the MRI session were selected if they had high IAcc. Thus, for this correlation there might be limited variance in the IAcc and Heartbeat Detection scores as we do not have individuals from the lower end of the spectrum on both scales.

Taken together, the behavioural performance between the two conditions was comparable although participants reported higher confidence for the Somatosensory condition. Here we interpret confidence ratings as subjective difficulty perceiving the stimuli and therefore infer that the Somatosensory Detection Task was subjectively perceived as easier.

### Focusing on cardiac and somatosensory signals

First, we looked at simple main effects of Cardiac and Somatosensory focus conditions (i.e. Cardiac Focus > baseline and Somatosensory > baseline). Both contrasts evoked a robust activation encompassing parietal, frontal and occipital areas (see Table 1 for details). Next, to study the extent of this overlap we conducted a formal conjunction analysis. The analysis confirmed a large overlap in the pattern of activation in these two conditions (Fig. 3A, Table 1). These include the right frontal operculum cortex extending towards insular cortex and inferior frontal gyrus, the lateral occipital cortex, bilaterally, extending towards angular gyrus and superior parietal cortex, fusiform gyrus, the supramarginal gyrus as well as juxtapositional lobule cortex (also known as supplementary motor area) extending into paracingulate cortex. Together these analyses show that cardiac and somatosensory focus recruit broadly the same regions.

**Table 1.** Results of the simple univariate analysis, looking at the focus to cardiac and somatosensory stimuli.

| Cluster Size<br>(Voxels) | P | Z-MAX | Coordinates |  |  | Side | Peak Activation Region |
| --- | --- | --- | --- | --- | --- | --- | --- |
|  |  |  | X | Y | Z |  |  |
| Cardiac Focus > Somatosensory Focus |  |  |  |  |  |  |  |
| 413 | < .001 | 4.2 | -20 | 22 | 56 | Left | Superior Frontal Gyrus |
| 400 | < .001 | 4.6 | 44 | -76 | 36 | Right | Lateral Occipital Cortex, superior division |
| 263 | .004 | 4.09 | 26 | 12 | 64 | Right | Superior Frontal Gyrus |
| 211 | .013 | 4.27 | -48 | -60 | 36 | Left | Lateral Occipital Cortex, superior division |
| Cardiac Focus > Baseline |  |  |  |  |  |  |  |
| 18567 | < .001 | 6.56 | 32 | 28 | 2 | Right | Frontal Orbital cortex |
| 13795 | < .001 | 5.64 | -58 | -46 | 16 | Left | Supramarginal Gyrus |
| 600 | < .001 | 5.49 | -34 | -90 | -10 | Left | Lateral Occipital cortex |
| 230 | .018 | 4.12 | 64 | -20 | 26 | Right | Supramarginal Gyrus |
| Conjunction (Cardiac Focus ∩ Somatosensory Focus) | | | | | | | |
| 37139 | < .001 | 6.24 | -6 | 10 | 56 | Left | Superior Frontal Gyrus |
| 2545 | .005 | 5.43 | 34 | -90 | -4 | Right | Lateral Occipital Cortex |
| Somatosensory Focus > Baseline |  |  |  |  |  |  |  |
| 17457 | < .001 | 6.27 | -8 | 10 | 54 | Left | Superior Frontal Gyrus |
| 3317 | < .001 | 6.02 | 62 | -22 | 20 | Right | Parietal Operculum Cortex |
| 1300 | < .001 | 5.57 | 34 | -90 | -2 | Right | Lateral Occipital Cortex |
| 959 | < .001 | 4.81 | 6 | -28 | 24 | Right | Cingulate gyrus, posterior division |
| 396 | .001 | 6.43 | -34 | -92 | -2 | Left | Occipital Pole |
| 389 | .001 | 4.18 | 18 | -12 | 10 | Right | Thalamus |

**Figure 3.**
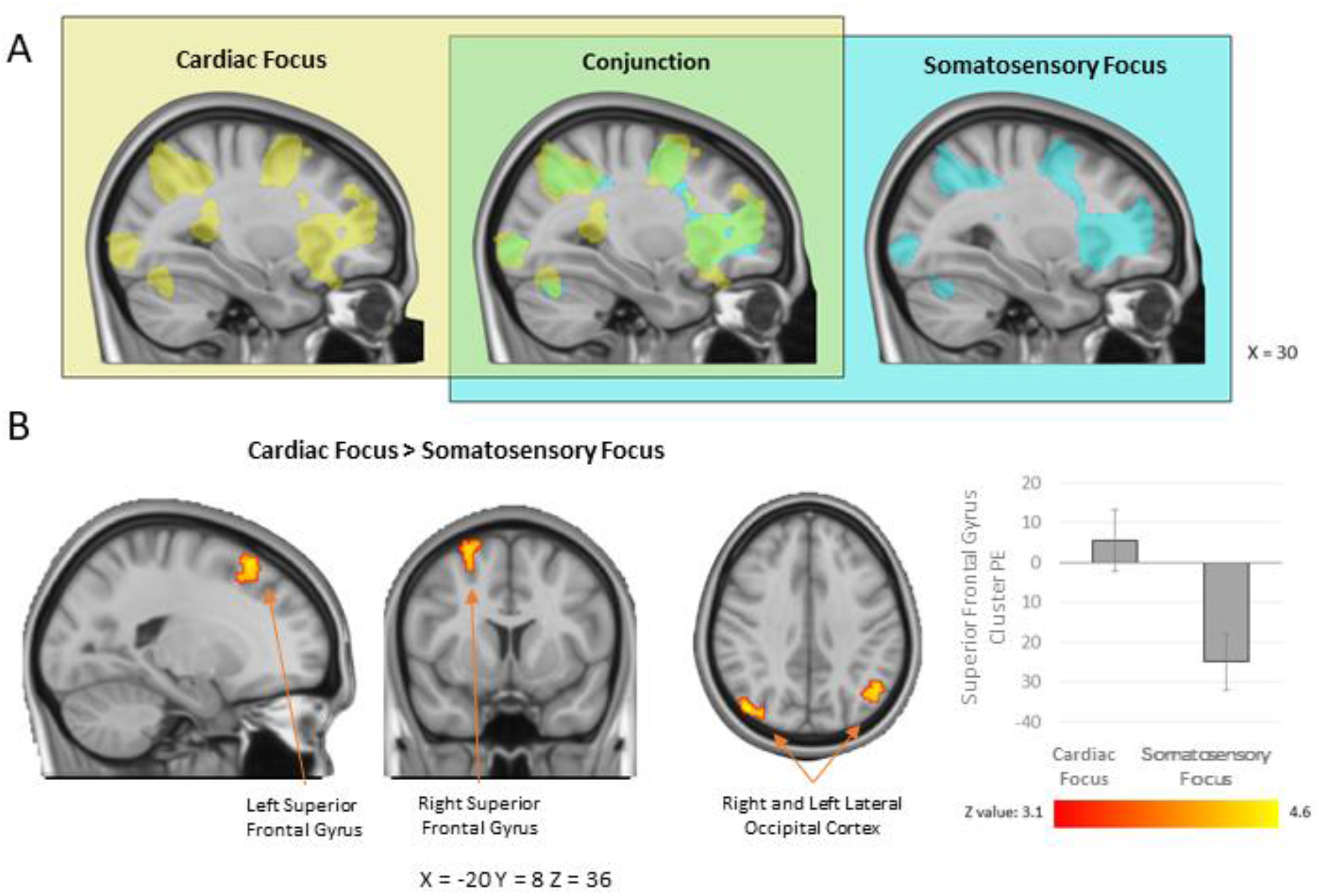
Results of the Univariate Analyses. (A) Regions activated during Cardiac Focus vs baseline (in yellow) and Somatosensory Focus condition vs baseline (in blue) and the results of the conjunction analysis between these two contrasts (in green). (B) Regions showing greater activation in the Cardiac Focus vs Somatosensory Focus condition. Bar plot represents the parameter estimates (PE) averaged over the whole cluster, error bars represent one standard error of the mean. All images are presented in the radiological convention: the right side of the brain is depicted in the left side of the image with coordinated in the MNI space.

In terms of differences between the focus conditions, that is depending on whether participants were instructed to focus on cardiac or somatosensory signals, the **Cardiac Focus > Somatosensory Focus** contrast yielded increased prefrontal (superior frontal and middle frontal gyri) as well as occipital (lateral occipital cortex extending into the angular gyrus) activation (Fig. 3B, Table 1). The reverse contrast **Somatosensory > Cardiac Focus** did not result in any suprathreshold clusters.

### Conscious perception of cardiac and somatosensory signals

We next investigated the neural correlates of consciously detected (Hits) and undetected (Misses) sensations across both conditions, as well as for each condition alone. For the detection by condition interaction effect [(Hits-Misses Cardiac) vs (Hits – Misses Somatosensory)], there were no suprathreshold clusters. Constricting the analysis to bilateral insular cortex (ROI analysis) also yielded no suprathreshold voxels. This suggests that detection of signals across both interoceptive and somatosensory domains engaged overlapping neural networks.

The main effect **Hits > Misses contrast** revealed a robust activation encompassing cortical (frontal, parietal and occipital) as well as subcortical areas bilaterally. These included precentral gyri, inferior, middle and superior frontal gyri, paracingulate cortex, insula, thalamus, putamen and caudate, brain stem, supramarginal gyrus, superior parietal lobule, postcentral gyri, lateral occipital cortex and precuneus (Fig 4A, Table 2). We followed this analysis with a formal conjunction analysis, looking at the brain areas that show overlapping activity when heartbeats and somatosensory stimuli were correctly detected. Indeed, we observed a robust overlap within all clusters (Fig 4B, Table 2). Nevertheless, the activation pattern for the Somatosensory condition seemed to be more widespread, particularly in the frontal and temporal areas, and also extending towards cerebellum.

**Table 2.** Results of the complex univariate analysis, investigating differences between consciously and non-consciously perceived sensations.

| Cluster Size<br>(Voxels) | P | Z-MAX | Coordinates |  |  | Side | Peak Activation Region |
| --- | --- | --- | --- | --- | --- | --- | --- |
|  |  |  | X | Y | Z |  |  |
| Main Effect: Hits > Misses |  |  |  |  |  |  |  |
| 23071 | < .001 | 6.00 | -10 | -14 | 6 | Left | Thalamus |
| 11050 | < .001 | 6.72 | 50 | -38 | 46 | Right | Supramarginal gyrus |
| 543 | .001 | 5.46 | 30 | -66 | -26 | Right | Cerebellum |
| 405 | .005 | 5.49 | -26 | -70 | -22 | Left | Occipital fusiform gyrus |
| 337 | .011 | 4.92 | 56 | -32 | -14 | Right | Inferior temporal gyrus |
| Hits > Misses Cardiac |  |  |  |  |  |  |  |
| 3008 | < .001 | 5.01 | 54 | -42 | 56 | Right | Supramarginal Gyrus |
| 2662 | < .001 | 4.79 | -48 | -46 | 56 | Left | Supramarginal Gyrus |
| 1579 | < .001 | 4.72 | 16 | -10 | 14 | Right | Thalamus |
| 1335 | < .001 | 4.69 | -56 | 10 | 40 | Left | Middle Frontal Gyrus |
| 823 | < .001 | 4.35 | 52 | 6 | 20 | Right | Precentral Gyrus |
| 485 | .003 | 4.24 | 26 | 0 | 50 | Right | Middle Frontal Gyrus |
| 275 | .032 | 3.92 | -34 | 2 | 64 | Left | Middle Frontal Gyrus |
| Hits > Misses Somatosensory |  |  |  |  |  |  |  |
| 17454 | < .001 | 6.81 | 48 | 16 | 28 | Right | Inferior Frontal Gyrus, pars opercularis |
| 9734 | < .001 | 6.53 | 44 | -42 | 44 | Right | Supramarginal Gyrus |
| 1232 | < .001 | 6.15 | -2 | 20 | 48 | Left | Paracingulate Gyrus |
| 350 | .004 | 5.19 | 28 | -68 | -26 | Right | Cerebellum |
| 320 | .006 | 4.69 | -26 | -70 | -24 | Left | Cerebellum |
| 317 | .006 | 4.69 | 56 | -32 | -14 | Right | Inferior Temporal Gyrus |
| Main Effect: Misses > Hits |  |  |  |  |  |  |  |
| 3387 | < .001 | 5.78 | 14 | -84 | 28 | Right | Cuneal cortex |
| 1845 | < .001 | 4.98 | 6 | 66 | -2 | Right | Frontal pole |
| 1004 | < .001 | 5.10 | -26 | -44 | -14 | Left | Temporal fusiform cortex |
| 909 | < .001 | 5.00 | 24 | -46 | -12 | Right | Lingual gyrus |
| 676 | < .001 | 5.62 | -48 | 0 | -22 | Left | Superior temporal gyrus |
| 274 | .026 | 4.48 | 38 | 12 | -26 | Right | Temporal pole |
| Misses > Hits Cardiac |  |  |  |  |  |  |  |
| 562 | .001 | 4.56 | 6 | 64 | -2 | Right | Frontal Pole |
| 447 | .004 | 3.93 | 8 | -48 | 32 | Right | Cingulate Gyrus, posterior division |
| Misses > Hits Somatosensory |  |  |  |  |  |  |  |
| 3111 | < .001 | 5.66 | 18 | -84 | 26 | Right | Lateral Occipital Cortex |
| 995 | < .001 | 5.03 | -26 | -42 | -14 | Left | Temporal fusiform Cortex |
| 967 | < .001 | 4.68 | 16 | 50 | 2 | Right | Paracingulate Gyrus |
| 838 | < .001 | 4.35 | 26 | -64 | -6 | Right | Occipital Fusiform Gyrus |
| 776 | < .001 | 5.58 | -50 | -2 | -24 | Left | Middle Temporal Gyrus |
| **Conjunction (Hits > Misses Cardiac ∩ Hits > Misses Somatosensory)** |  |  |  |  |  |  |  |
| 2692 | < .001 | 5.01 | 54 | -42 | 56 | Right | Supramarginal Gyrus |
| 2414 | < .001 | 4.79 | -48 | -46 | 56 | Left | Supramarginal Gyrus |
| 1227 | < .001 | 4.13 | -18 | 20 | 2 | Left | Caudate |
| 1044 | .001 | 4.21 | 22 | 10 | 8 | Right | Putamen |
| 960 | .002 | 4.69 | -56 | 10 | 40 | Left | Middle Frontal Gyrus |
| 662 | .009 | 4.35 | 52 | 6 | 20 | Right | Precentral Gyrus |
| 450 | .034 | 4.24 | 26 | 0 | 50 | Right | Middle Frontal Gyrus |

**Figure 4.**
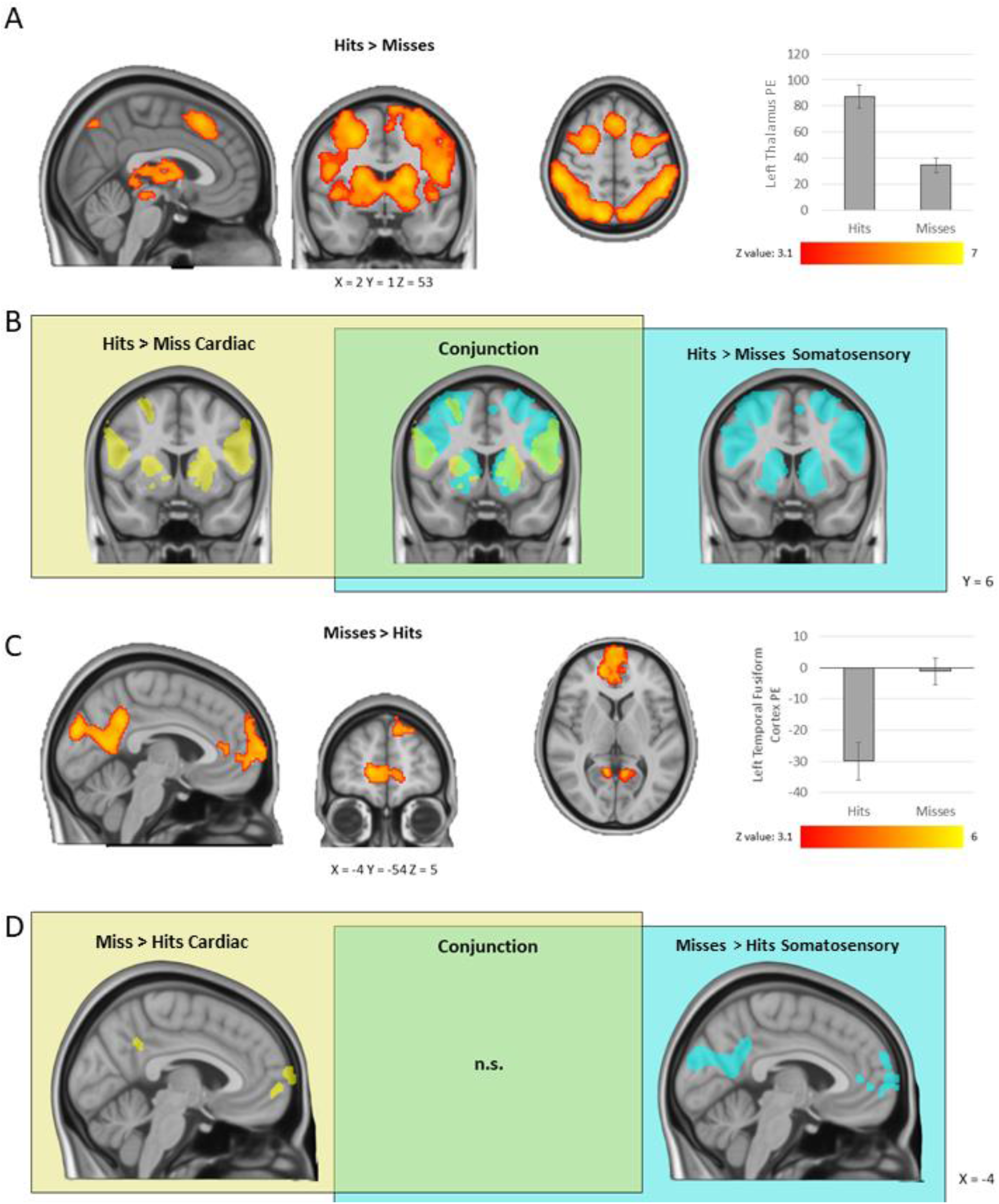
Results of the complex univariate analysis, investigating differences between consciously and non-consciously perceived sensations. Main effects analysis of Hits > Misses (A) and the conjunction analysis results (B) showing areas of greater activation during Hits vs Misses for each focus condition and the results of the conjunction analysis (in green). Main effect analysis of Misses > Hits (C) and the activations for each condition separately (D). All images are presented in the radiological convention: the right side of the brain is depicted in the left side of the image with coordinated in the MNI space. Bar plots represent the parameter estimates (PE) averaged over the whole cluster, error bars represent one standard error of the mean.

The **reverse main effects contrast (Misses > Hits)** revealed activations in bilateral temporal fusiform cortex, lingual gyrus, hippocampus and parahippocampal gyrus, inferior and middle temporal gyri, precuneus cortex, cingulate gyrus, fusiform gyrus, cuneal cortex as well as lateral occipital cortex and lingual gyrus (Fig 4C, Table 2). The conjunction analysis revealed no significant overlap of processing missed sensations of both types of sensations (Fig 4D). For the Cardiac condition, the activation was limited to frontal pole and posterior cingulate gyrus, extending towards precuneus. The activation for the Somatosensory condition also encompassed lateral occipital cortex, temporal cortex, hippocampus and parahippocampal gyrus, cueneal and precuneus cortex.

### Psycho-physiological interactions

We used the gPPI to test the hypothesis that the functional connectivity strength of the right insula cortex ROI would be differentially modulated by the conscious detection (i.e. Hits > Misses) of Cardiac versus Somatosensory stimuli. Indeed, we observed a significant interaction effect whereby the functional connectivity of the right insula ROI was greater for consciously detected heartbeats than somatosensory stimuli (Table 3, Fig 5). Specifically, conscious detection of heartbeats was related to increased connectivity with the lateral occipital cortex extending towards cuneal and precuneus cortex, right middle temporal gyrus, lingual gyrus, occipital pole, left supramarginal gyrus extending towards postcentral gyrus as well as left planum temporale extending towards parietal and central operculum cortex. These differences suggest that top-down attentional processes and conscious detection of different sensory events might modulate the right insular cortex functional connectivity. As a follow-up, we repeated the gPPI analysis separately for the Cardiac and Somatosensory conditions revealing that the interaction was primarily driven by significant changes in right insular functional connectivity during heartbeat perception. No significant changes in connectivity were found for the somatosensory condition (see Supplementary Materials for details).

**Table 3.**
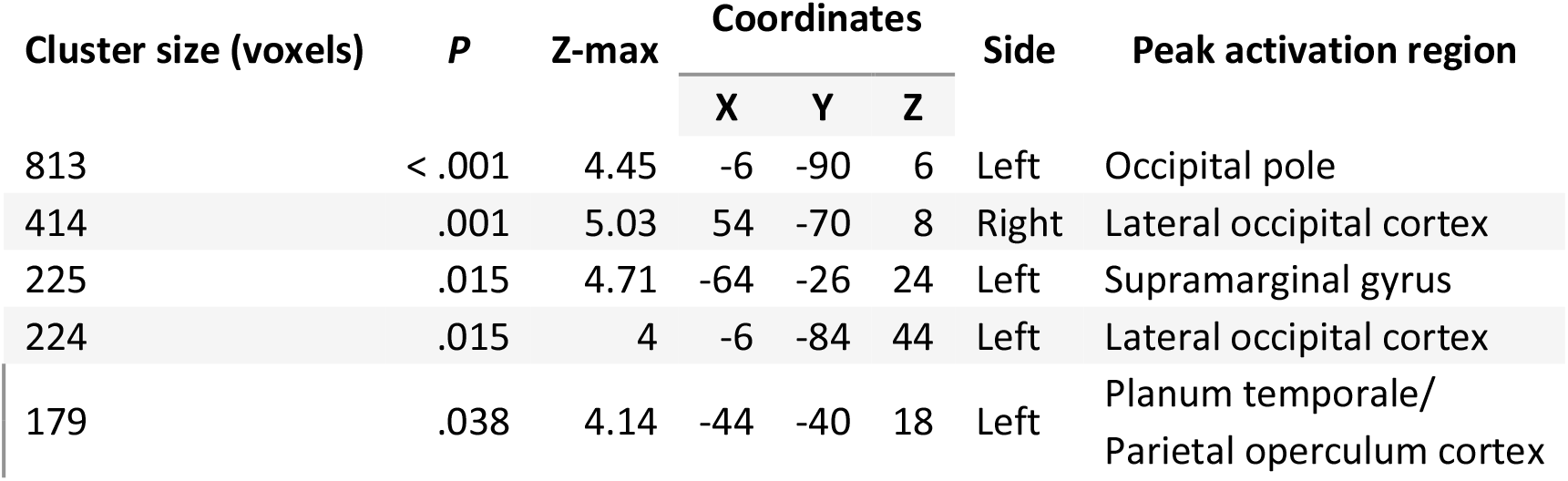
PPI results for Cardiac Focus vs Somatosensory Focus during Hits relative to Misses contrast. The coordinates for clusters maxima are presented in MNI space.

| Cluster size (voxels) | P | Z-max | Coordinates |  |  | Side | Peak activation region |
| --- | --- | --- | --- | --- | --- | --- | --- |
|  |  |  | X | Y | Z |  |  |
| 813 | < .001 | 4.45 | -6 | -90 | 6 | Left | Occipital pole |
| 414 | .001 | 5.03 | 54 | -70 | 8 | Right | Lateral occipital cortex |
| 225 | .015 | 4.71 | -64 | -26 | 24 | Left | Supramarginal gyrus |
| 224 | .015 | 4 | -6 | -84 | 44 | Left | Lateral occipital cortex |
| 179 | .038 | 4.14 | -44 | -40 | 18 | Left | Planum temporale/<br>Parietal operculum cortex |

**Figure 5.**
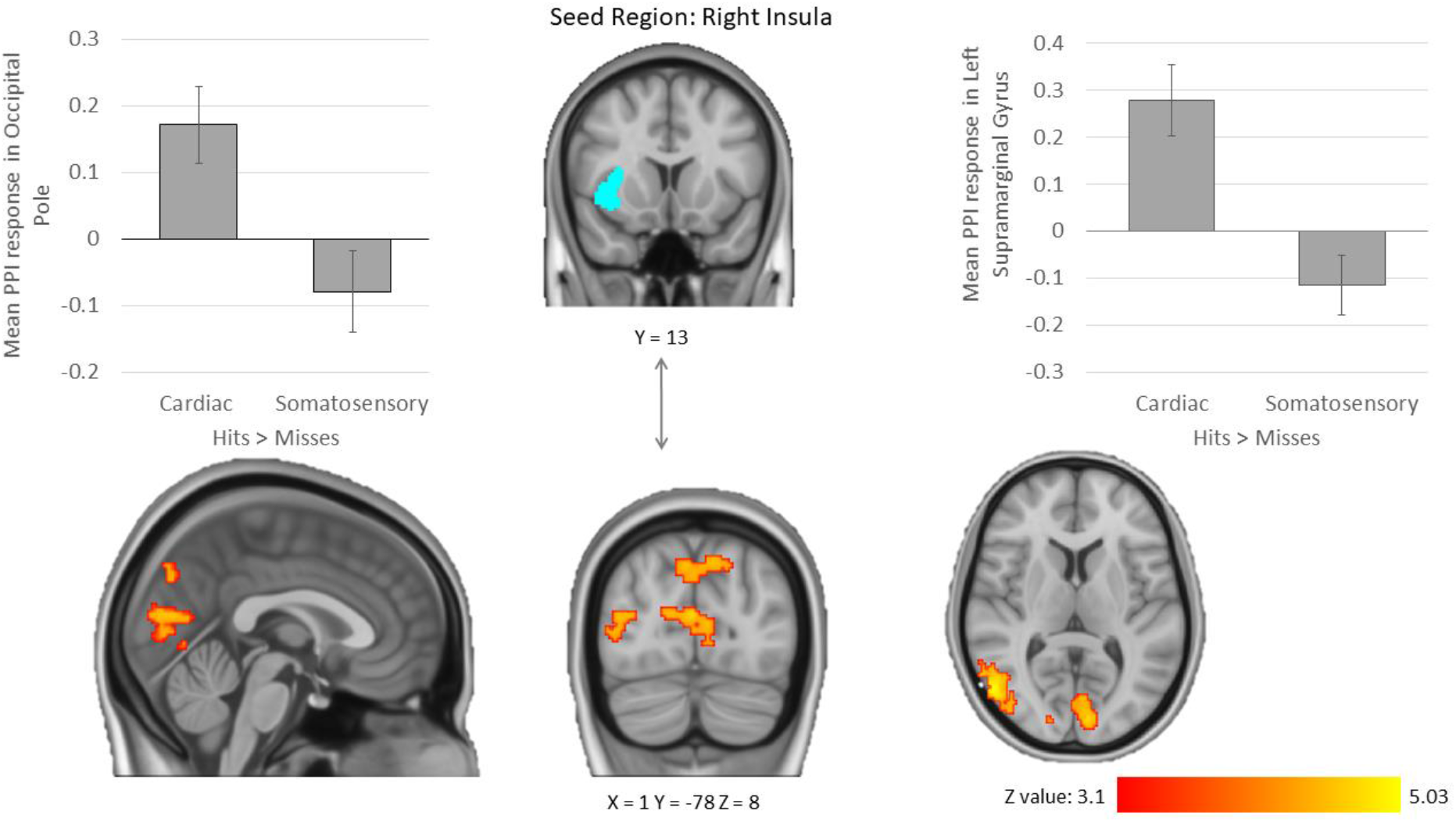
PPI results showing greater functional connectivity between the right insula seed and occipital and parietal areas in the Cardiac Focus vs Somatosensory Focus during Hits relative to Misses contrast. Images are presented in the radiological convention: the right side of the brain is depicted in the left side of the image with coordinated in the MNI space. Bar plots represent the PPI response averaged across the whole cluster; error bars represent one standard error of the mean.

## Discussion

The current study used a novel Heartbeat-Somatosensory detection paradigm to better understand the neural correlates of attention to interoceptive and somatosensory stimuli and their conscious detection. Additionally, we investigated the neural networks underpinning conscious and non-conscious perception of these stimuli. Overall, we observed a robust overlap in the pattern of activation evoked by both Focus conditions in frontal, parietal and occipital areas, including insular cortex. Correct detection of stimuli (Hits > Misses), heartbeats and somatosensory stimuli alike, evoked greater activation in frontal, parietal occipital, and insular cortex areas, as well as subcortical areas and brain stem. On the other hand, undetected stimuli (Misses > Hits) evoked greater activations in frontal pole, posterior cingulate and precuneus as well as temporal areas. Nevertheless, we also observed some important differences. Cardiac Focus yielded increased prefrontal (superior frontal and middle frontal gyri) and occipito-parietal (lateral occipital cortex extending into angular gyrus) activation relative to the Somatosensory Focus condition. Additionally, psychophysiological interactions analysis revealed that right insular cortex functional connectivity was modulated by the conscious detection of interoceptive and exteroceptive sensations differently, showing greater connectivity with a set of occipito-parietal regions during Cardiac compared to Somatosensory Focus. The subsequent analysis further revealed that this interaction was driven by the altered anterior insula connectivity mainly during the cardiac condition. Together, our results suggest a large degree of convergence in neural correlates underlying attention to and conscious detection of interoceptive and (proximal) exteroceptive stimuli.

### Cardiac versus somatosensory focus

Focus to interoceptive signals (Cardiac Focus condition) yielded increased prefrontal (superior frontal and middle frontal gyri) as well as occipital (lateral occipital cortex extending into the angular gyrus) activation compared to Somatosensory Focus condition. Both, prefrontal and occipital activations in interoceptive conditions have been identified previously (Critchley et al., 2004; Stern et al., 2017; Wang et al., 2019). Increased activation in visual areas may reflect higher visual attention and increased engagement in mental imagery necessary to integrate perceived heartbeats with corresponding visual stimuli (colours), particularly in the cardiac condition where this task is more difficult. Moreover, for the somatosensory condition, the stimulation was always applied to the same location on the skin; whereas, participants could focus on different body parts in the cardiac condition to detect their heartbeat, which may have differed across trials and could have relied on greater mental imagery in the cardiac condition. The superior and middle frontal gyri are both strongly involved in attentional and cognitive control in general (Bauer, Barrios, & Díaz, 2014; Talati & Hirsch, 2005; Weber & Huettel, 2008; Wilbertz et al., 2014), particularly in focused attention tasks and meditation (Brefczynski-Lewis, Lutz, Schaefer, Levinson, & Davidson, 2007; Doll et al., 2016). For example, the left superior frontal gyrus/middle frontal gyrus area consistently showed increased activation in expert meditators during focused attention meditation (Brefczynski-Lewis et al., 2007). Therefore, enhanced activity in these areas may reflect higher cognitive and attentional resources engaged in task performance during Cardiac Focus vs Somatosensory Focus Condition. These results are consistent with behavioural findings, whereby participants showed lower confidence in the Cardiac than Somatosensory condition, suggestive of the former being subjectively more difficult. At the same time, confidence ratings should be cautiously interpreted. Confidence ratings can be modulated by objective task difficulty (Whitmarsh, Oostenveld, Almeida, & Lundqvist, 2017) but also other factors such as general metacognitive abilities (Fleming & Lau, 2014) or individual differences in confidence independent of task difficulty (Beck, Peña-Vivas, Fleming, & Haggard, 2019). For example, regarding somatosensory stimulation, Grund et al. (2021) have recently shown that participants report lower confidence ratings for near-threshold hits compared to near-threshold misses, despite the same intensity (near-detection threshold) and hence the same objective difficulty. Similarly, elevated occipital activation may reflect increased visual attention. The angular gyrus is considered to be a cross-modal integrative hub for converging information from different sensory modalities (for review see Seghier, 2013). Given the relatively higher perceived difficulty of our Heartbeat Detection task, which involves integration of visual cues with internal bodily signals, the angular gyrus involvement as an integrative hub seems key.

However, we did not find any differences in activation between the Cardiac and Somatosensory focus conditions within the insula or the anterior cingulate cortex, regions commonly considered to be the key elements of interoceptive processing (A. D. (Bud) Craig, 2009a; Critchley et al., 2004; Salvato et al., 2019; Schulz, 2016). Importantly though, the role of insula extends well beyond interoception and encompasses salience processing (Uddin, 2015), emotional awareness and regulation (Critchley, 2009; Phan, Wager, Taylor, & Liberzon, 2002; Shafritz, Collins, & Blumberg, 2006), as well as sensory processing and multimodal integration more generally (Avery et al., 2015; Plailly, Radnovich, Sabri, Royet, & Kareken, 2007; Simmons et al., 2013; Y. Suzuki et al., 2001). Indeed, previous neuroimaging studies showed that vibrotactile stimulation using pneumatic devices, as in the present study, predominantly elicits activation of the primary and secondary somatosensory cortex as well as the insula and the thalamus (e.g., Briggs et al., 2004; Chakravarty et al., 2009; Chang et al., 2009; Golaszewski et al., 2006; Nelson et al., 2004). These regions show overlap with the network we identified by conjunction analysis of Cardiac and Somatosensory Focus conditions in the current study.

Overall, the focus to cardiac signals and somatosensory stimuli in our study showed highly overlapping activation patterns in several brain regions, including the insula, the cingulate, frontal gyri, somatomotor and occipital regions. This network of activity is highly congruent with the anatomical structures of the interoceptive network identified in previous studies (e.g., Critchley et al., 2004; Kuehn et al., 2016; Pollatos et al., 2007a; Stern et al., 2017; Zaki, Davis, & Ochsner, 2012b). The extent of overlap revealed in the conjunction analysis points to a large degree of commonality between the two modalities of body processing. Such large overlap may indicate an important role of these structures for bodily self-consciousness but also suggests that somatosensory pathways, rather than solely interoceptive pathways, participate in cardioception (Khalsa, Rudrauf, Feinstein, & Tranel, 2009).

The overlap was found in several parietal regions, such as supramarginal gyrus (SMG), angular gyrus, and superior parietal lobule, all of which are implicated in multisensory processing and integration. A recent meta-analysis revealed that the internal (interoceptive) and external (related to the experience of body-ownership) signals integration occurs in the SMG bilaterally together with a right-lateralized set of areas such as the precentral, postcentral, and superior temporal gyri (Salvato et al., 2019). These higher-order brain areas are involved in integrating multisensory signals, and in recalibrating information from different incoming channels and spatial frames of reference (Salvato et al., 2019). The right SMG is also important for proprioception (Ben-Shabat, Matyas, Pell, Brodtmann, & Carey, 2015), while left SMG is associated with decoding of self-location (Guterstam, Björnsdotter, Gentile, & Ehrsson, 2015) and perceiving limbs in space in a body-centred reference (Brozzoli, Gentile, & Henrik Ehrsson, 2012). It has been suggested that primary somatosensory areas together with left fronto-parietal areas are involved in processing proprioceptive and interoceptive bodily information that underlies body-representations (Bauer, Díaz, Concha, & Barrios, 2014).

We also found an extensive overlap in activation in the lateral occipital cortex. Prior research identified regions of lateral occipito-temporal cortex (extrastriate body area and the fusiform body area) to be involved in body processing, not only when viewing images of the human body and body parts (Costantini, Urgesi, Galati, Romani, & Aglioti, 2011; Taylor, Wiggett, & Downing, 2007; Urgesi, Candidi, Ionta, & Aglioti, 2007), but also when engaging in mental imagery of embodied self-location (Arzy et al., 2006), mental manipulation of body parts (Kikuchi et al., 2017) as well as experiencing illusory body ownership (Limanowski, Lutti, & Blankenburg, 2014). Possibly, while focusing on perception of one’s heartbeat or on detecting stimuli applied to one’s hand, participants saw the relevant body parts in their minds’ eye.

Overall, our results point to a large degree of convergence in neural mechanisms underlying attentional mechanism directed towards interoceptive (heartbeats) and exteroceptive (vibrotactile) stimuli. We found little evidence for divergence between these two processes. To some extent, these results may reflect our design, namely the types of stimuli used (proximal, vibrotactile stimulation), their continuing presence throughout and the relative difficulty of the task, but also the inherent convergence of bodily-related signals. Our brains may be primarily wired to integrate rather than separate proximal exteroceptive and interoceptive bodily signals.

### Conscious and non-conscious stimuli detection

Apart from the main and conjunctive effects of attention directed internally or externally, we also investigated the aspects of conscious perception of stimuli. We did not find any interaction effect regarding detection accuracy (felt vs missed sensations) and focus condition. This may reflect high task-demands and comparable difficulty of the tasks, as determined by behavioural performance that was found to be correlated between the two conditions. Moreover, in order to match the conditions as closely as possible, we ensured there was a train of somatosensory stimuli throughout the cardiac focus blocks. This was important to mimic the continuous presence of the heart beat during the somatosensory blocks, but likely increased the difficulty of the task and reduced our ability to detect differences in the BOLD response between the conditions. Instead, correctly detected sensations compared to missed sensations (Hits > Misses) across both conditions evoked activations in frontal (inferior, middle and superior frontal gyri, paracingulate cortex), somatomotor areas, the insula, as well as subcortical areas (thalamus, putamen and caudate), brain stem, SMG, superior parietal lobule, lateral occipital cortex, and precuneus. This pattern of activation was highly consistent across both conditions as revealed by the conjunction analysis. This pattern of activation bares resemblance to the salience network and executive control network (Seeley et al., 2007). The salience network consists of anterior cingulate cortex and orbital frontal insula; both regions co-activate in response to varied forms of salience (Seeley et al., 2007). Moreover, as a part of this network, anterior insula is considered an integral hub enabling dynamic switches between externally and internally oriented attention (Menon & Uddin, 2010; Uddin, 2015). The executive control network encompasses dorsolateral prefrontal and parietal cortices and is thought to underlie many goal-directed processes such as sustained attention and working memory as well as response selection and suppression (Seeley et al., 2007). Therefore, given the role of these networks in detecting salience and goal-directed attentional switches, the activation of these regions in consciously detected bodily/external cues is not surprising.

In contrast, the reversed comparison, Misses > Hits, evoked no significantly overlapping areas of activation across both conditions. Missed heartbeats were associated with frontal pole, posterior cingulate and precuneus activation, while missed Somatosensory stimuli were also associated with more widespread activation in frontal and temporal regions. These results suggest some degree of separation between un-conscious processing of cardiac and somatosensory stimuli. Nevertheless, the main effect of Misses > Hits across both conditions evoked frontal pole, posterior cingulate and precuneus as well as temporal activations. Overall, these activations show some resemblance to the default mode network (DMN) which encompasses the precuneus/cingulate cortex, medial prefrontal cortex as well as areas of parietal cortex (Mason et al., 2007; Raichle et al., 2001). The DMN shows lower activation during task relative to resting condition. Nevertheless, it is thought to play a far more important role than just allowing us to daydream, as it is linked to self-referential activity, reflecting upon one’s own mental state, introspection and autobiographical memory (Andrews-Hanna, Smallwood, & Spreng, 2014; D’Argembeau et al., 2005; Gusnard & Raichle, 2001). Therefore, the greater activation of the DMN during missed trials, may reflect simple off-task activity (inattention), but it could also reflect aspects of self-reflection. This clear differentiation between task-positive networks, underlying aspect of attentional control and salience processing during correct detections and greater activation of task-negative DMN during missed trials may determine performance in the task.

Our findings differ from previous studies looking at conscious detection of exteroceptive stimuli (Meneguzzo et al., 2014). Indeed, this meta-analysis highlighted greater activity in left anterior cingulate cortex and mid-caudal anterior cingulate cortex with conscious detection of stimuli, which was the opposite to the one reported here. However, this discrepancy may be driven by the nature of the stimuli measured: in their meta-analysis only exteroceptive, visual and tactile (rectal stimulation in clinical population), stimuli were considered. Previous studies measuring attentional fluctuations to vibrotactile stimuli have been associated with increased parietal activity (Schmidt & Blankenburg, 2018), which is in line with the associations for cardiac and somatosensory perception in the current study. This highlights how the modality being measured can impact the pattern of the BOLD response across the brain, however, importantly, this was not the case for the cardiac and somatosensory stimuli in the current study. Goltz et al., (2015) found that connectivity with the intraparietal sulcus was associated with attentional fluctuations in vibrotactile perception. Therefore, although the regions recruited when focusing on cardiac and somatosensory stimuli converge, the network dynamics between these regions may differentiate perception across these modalities.

### Right insula task-related functional connectivity changes

Even though we did not find a focus condition by detection interaction, the right insula functional connectivity showed an interaction effect. Specifically, conscious detection of heartbeats (Hits > Misses) was related to greater functional connectivity between the right insula ROI and areas encompassing occipital (lateral occipital cortex, lingual gyrus, occipital pole), parietal (cuneal and precuneus cortex, left SMG extending towards postcentral gyrus, parietal and central operculum cortex) as well as temporal cortices (right middle temporal gyrus, left planum temporale), relative to the conscious detection of somatosensory stimuli. Interestingly, the right insula connectivity was associated with the detection of cardiac stimuli only. Therefore, conscious detection of heartbeats was related to higher degree of communication between the right insula, the area considered a key hub of interoceptive processing (A. D. (Bud) Craig, 2003, 2009a; Critchley et al., 2004), and other areas of the interoceptive network (i.e. postcentral gyrus, secondary somatosensory cortex) and as well as the set of regions associated with body self-ownership (occiptotemporal and parietal areas) (Salvato et al., 2019). Noteworthy, our results indicate that conscious perception of heartbeats is related to greater functional connectivity of the right anterior insula and SMG, the cortical region where the processing of both body ownership and interoception converges (Salvato et al., 2019). The increased connectivity of insular ROI with the occipital cortex could be part of the long-term representation of the body involving its pictorial appearance and visualization (Bauer, Díaz, et al., 2014). Together, our results suggest that top-down attentional processes and conscious detection of different sensory events modulate the right insular cortex functional connectivity. Additionally, conscious perception of heartbeats was related to greater functional connectivity of the right insula and somatosensory cortices. Functional neuroimaging findings implicate insula and anterior cingulate cortices together with somatosensory regions in interoceptive awareness (Cameron & Minoshima, 2002; Critchley et al., 2004; Pollatos et al., 2007). Moreover, insula lesion research indicated that heart rate awareness was mediated by both somatosensory afferents from the skin and a network that included the insula and anterior cingulate cortex, suggesting that both of these pathways enable the perception of cardiac signals and states (Khalsa et al., 2009). Our results further suggest that insular and somatosensory cortices work together to form a conscious cardiovascular state detection.

Anterior insula activity is consistently activated in studies that elicit changes in autonomic arousal (Cameron & Minoshima, 2002; Critchley, 2002; Critchley, Corfield, Chandler, Mathias, & Dolan, 2000; Critchley, Mathias, & Dolan, 2001, 2002; Critchley et al., 2003). It is also activated by visceral stimulation (Aziz, Schnitzler, & Enck, 2000), olfactory and gustatory stimuli (Rolls, 2015; Smejkal, Druga, & Tintera, 2003), pain (Peyron et al., 2002), temperature (A. D. Craig, Chen, Bandy, & Reiman, 2000; Stern et al., 2017) and emotional processing (Wicker et al., 2003; Zaki et al., 2012). Right insula cortex activity is also enhanced in appraisal of emotions and bodily physiological state, suggesting that anterior insula serves as an interface between physiologically driven internal motivational states, emotional awareness and interpersonal behaviour (Terasawa, Shibata, Moriguchi, & Umeda, 2013). Together, this supports the notion that the right anterior insula, as playing a central role in interoceptive processes and representation of bodily arousal, engenders human awareness providing a substrate for subjective feeling states (A. D. (Bud) Craig, 2003, 2009a; Critchley et al., 2004).

Some limitations merit comment. As much as we made every effort to match both focus conditions as closely as possible, the somatosensory stimuli were present more frequently than heartbeats, due to subject’s heart rate’s decreasing throughout the duration of the task. One could argue that the occurrence of more somatosensory than cardiac events is a confound that could affect people’s performance. Yet, as we show above if anything people’s accuracy was similar, if not slightly better, for cardiac than somatosensory events. Recording ECG within an MRI scanner is extremely difficult, therefore although attempts were made to match the presentation rate of the tactile stimuli to that of the subject’s heartbeat during data collection, we were not able to measure heart rate in real time for the majority of subjects. The timing of each cardiac R-peak was determined after the scanning session following post-processing of the ECG signal. The Somatosensory Focus condition was also associated with higher confidence ratings than Cardiac Focus condition. However, given the lack of many differences between conditions it is unlikely that these differences were driving the results. Moreover, as the epoch duration (window of time during which participants could expect to feel the stimulus) was quite long relative to the average heartbeat cycle, both stimuli were present on the vast majority of the epochs and for some blocks, participants’ heart rate exceeded 80bpm. However, as this was very rare (happened only seven times across all blocks for all participants), therefore we do not think this affected our main analysis. The fast presentation rate also caused that there were some between-participant differences in the stimuli presentation frequency with some having no false alarms or correct rejections dependent on heart rate. This is a common problem with attempts to use signal detection theory to measure cardiac detection; it is difficult to ensure there are trials in which the heartbeat is absent particularly when a subject has a fast heart rate. Finally, we deliberately selected individuals who presented relatively good performance in our heartbeat detection task. We cannot exclude the possibility that individuals with significantly lower or higher interoceptive accuracy potentially may process sensory information coming from within and outside of the body in different ways.

### Summary and Conclusions

In line with our hypothesis, we found overlapping but dissociable activation patterns associated with both internally- (heartbeats) and externally- (somatosensation) oriented attention. The robust overlap included key areas typically associated with interoceptive processing, including insula, somatomotor cortices, cingulate cortex, suggesting their broader role in processing body-related information to construct and maintain body self-consciousness. Nevertheless, Cardiac Focus additionally evoked higher frontal and occipito-parietal areas in regions associated with cognitive control and multimodal integration. Importantly, this task provides an important advance towards experimental designs that move away from measuring interoceptive attention only to begin to delineate the neural correlates of conscious detection of interoceptive stimuli from other modalities. The correct detection of interoceptive and somatosensory sensations evoked overlapping activations in salience – control network, while missed sensations evoked activations in areas linked to the DMN. Although we did not observe an interaction with the conscious detection condition our gPPI analysis revealed that functional connectivity with the right insular cortex, a central hub for interoceptive processing, was modulated by conscious detection of heartbeats between focus conditions suggesting the role of top-down processes influencing insular connectivity. Due to the crucial role of multimodal information, including interoceptive, somatosensory, and proprioceptive information, in body-representation and awareness, these findings extend previous knowledge regarding the neural correlates of directed attention to internal and somatosensory stimuli and conscious as well as non-conscious processing of these sensations.

## Author contributions

C.P., R.A., and M.T. conceived and designed the experiments. C.P. carried out the experiments and performed ECG data preprocessing. A.H. and C.P. contributed to the behavioural data analysis. A.H. performed the fMRI data analysis. A.H. took the lead in writing the manuscript. All authors provided critical feedback and helped shape the research, analysis and the final version of the manuscript.

