## Supplementary Materials for "Neural divergence and convergence for attention to and detection of interoceptive and somatosensory stimuli"

### Behavioural performance

For completeness, in addition to Figure 2 in the main text, SFigure 1 presents the behavioural performance on the Heartbeat and Somatosensory Detection Task, whereby performance is presented as per signal detection theory approach. The figure clearly shows, that the correct responses were highly comparable across blocks and conditions. The number of misses and false alarms differed between conditions. This might be related to the fact that, overall, more somatosensory stimuli were present than heartbeats (see main text for details) and possibly participants were trying to 'match' the number of their responses between conditions.

Additionally, to check whether the performance differed across epochs (colourful crosses within each trial) not per block, we conducted an additional analysis whereby we collapsed correct detections (hits) and incorrect rejections (misses) per epoch for cardiac and somatosensory condition separately (SFigure 2). Repeated-measures analysis of variance (rm-ANOVA) indicated that for hits, there was a main effect of epoch [$F(2, 54) = 18.43$, $p < .001$, $\eta_p^2 = 0.41$] with significantly fewer correct detections during the first epoch relative to the second ($p < .001$) and third ($p = .014$) epochs. There was no significant main effect of condition [$F(1,27) = 0.55$, $p = .466$, $\eta_p^2 = 0.02$] but there was a significant condition x epoch interaction [$F(2, 54) = 5.20$, $p = .009$, $\eta_p^2 = 0.16$], with higher proportion of hits for somatosensory than cardiac condition during the first epoch, and fewer hits for somatosensory than cardiac condition during the third epoch. Regarding misses, there was also a significant main effect of epoch [$F(2, 54) = 39.55$, $p < .001$, $\eta_p^2 = 0.59$] with significantly more missed stimuli during the first epoch relative to the second ($p < .001$) and third ($p < .001$) epochs. There was also a significant main effect of condition [$F(1,27) = 12.65$, $p = .001$, $\eta_p^2 = 0.32$] with overall more misses for the somatosensory than cardiac condition. This likely reflects the fact that overall, more somatosensory stimuli were present than heartbeats (see main text for details). The condition x epoch interaction was insignificant [$F(2, 54) = 2.12$, $p = .130$, $\eta_p^2 = 0.07$]. Overall, these results suggest that within trial, the detection of stimuli, regardless of the condition, was the poorest for the first epoch but remained stable for the last two epochs. This probably reflects the fact that the inter-trial-interval was jittered and varied between 4 and 8s and participants were not prompted to re-engage with stimuli monitoring.

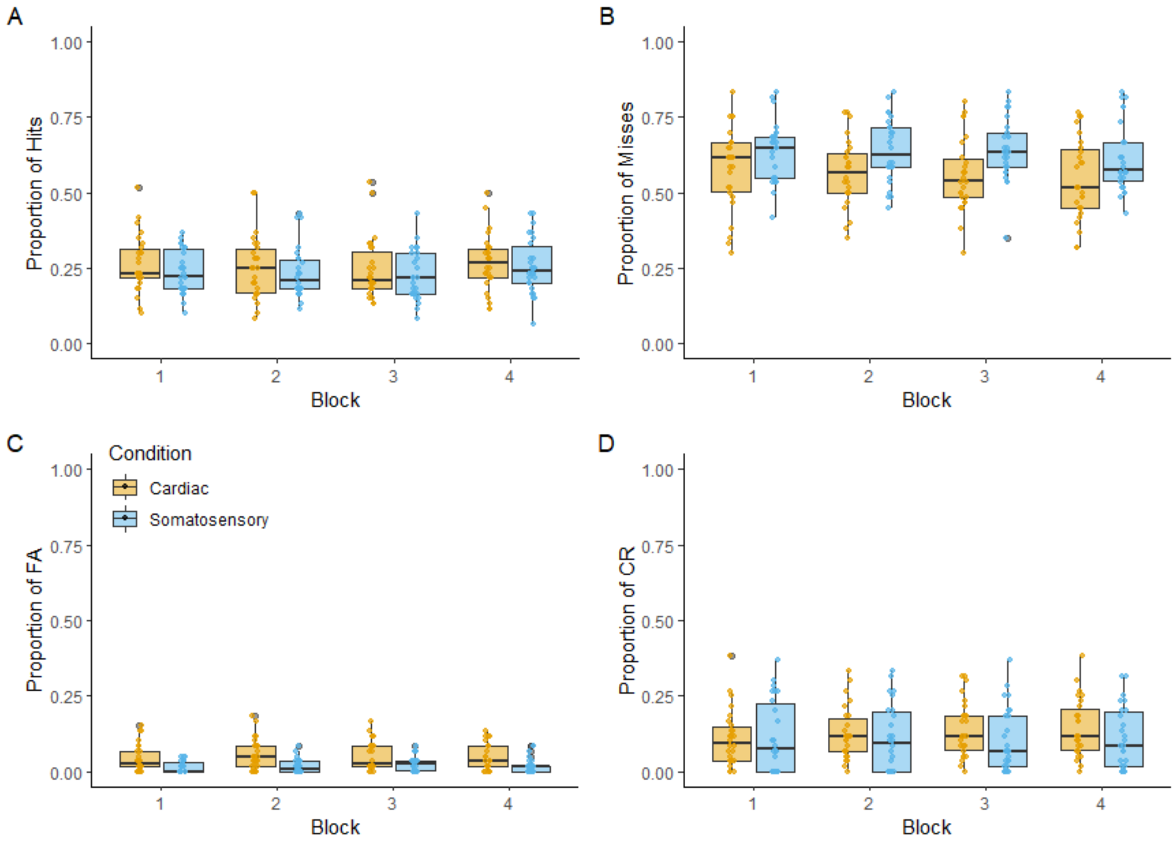


*SFigure 1 Performance on the Heartbeat/Somatosensory Detection Task with responses marked as per signal detection theory presented as a function of block and condition. A) Proportion of hits (correct detections). B) Proportion of misses (incorrect rejections). C) Proportion of false alarms (FA; incorrect detections). D) Proportion of correct rejections (CR).*

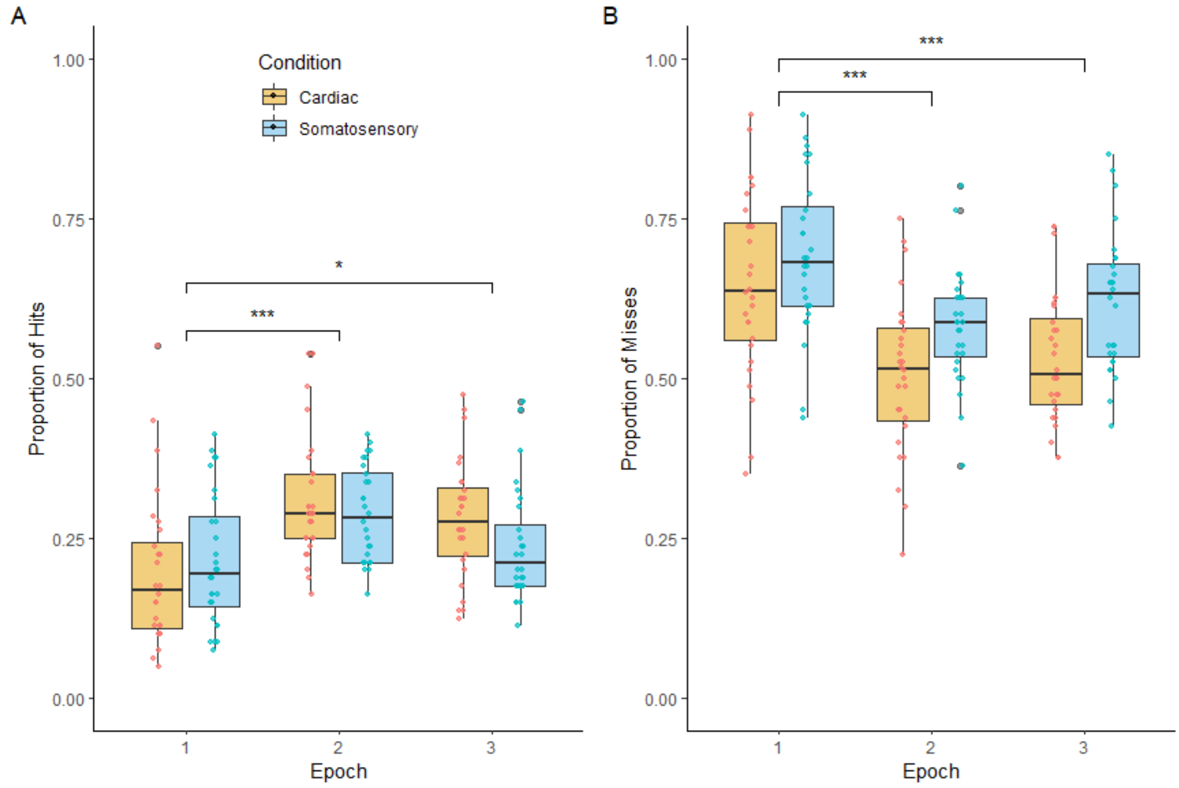


*SFigure 2 Proportion of (A) hits and (B) misses per epoch (each colourful cross) and condition. The epoch number corresponds to the colourful cross (i.e. red, green, blue) as presented to participants in the Heartbeat/Somatosensory Detection Task. *p < 0.05, ***p < 0.001*

### Follow-up PPI analysis

To follow-up on the PPI results, we repeated the PPI analysis separately for each focus condition (i.e. hits vs misses for Cardiac condition and hits vs misses for Somatosensory condition). Following the same methodology as described in the main text, we found that the right insula ROI showed higher functional connectivity for hits vs misses with the right supracalcarine cortex and left supramarginal gyrus extending into parietal operculum (SFigure 3, STable1). There were no suprathreshold results for the Somatosensory conditions and the conjunction was also insignificant. Thus, the significant interaction effect is mainly driven by changes in the right insular functional connectivity in the cardiac condition.

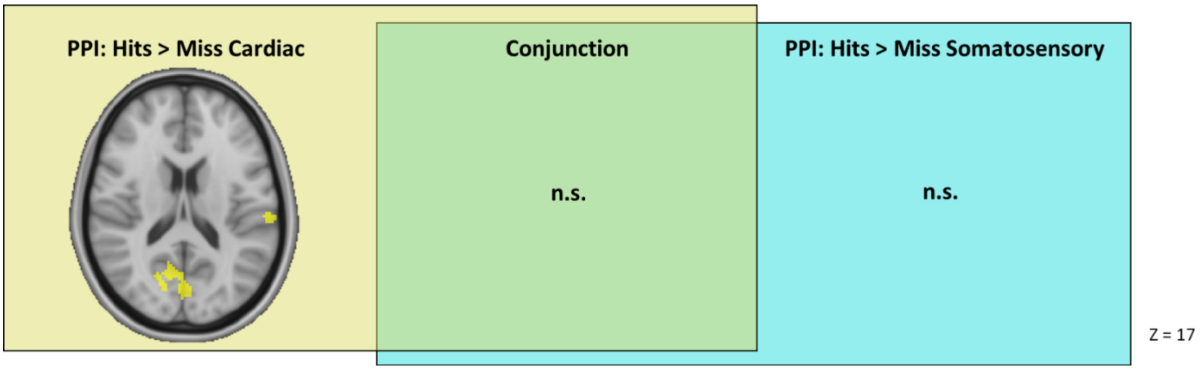


*SFigure 3 Results of the follow-up PPI analysis, investigating differences in right insular ROI functional connectivity between consciously and non-consciously perceived sensations for each focus condition separately and the results of the conjunction analysis (in green).*

*STable 1 PPI results for Cardiac Focus during Hits relative to Misses contrast. The coordinates for clusters maxima are presented in MNI space.*

| Cluster Size (Voxels) | P | Z-MAX | Coordinates | | | Side | Peak Activation Region |
|---|---|---|---|---|---|---|---|
| | | | X | Y | Z | | |
| **214** | .011 | 4.16 | 2 | -76 | 18 | Right | Supracalcarine cortex |
| **167** | .032 | 4.32 | -64 | -26 | 24 | Left | Supramarginal gyrus |

### Detailes of additional analysis

As a complementary approach, we conducted an additional analysis whereby we only modelled the onsets with respect to the heart beat or the somatosensory stimulus to specifically isolate the activation during detection of the stimulus as opposed to the whole epoch. This approach could provide a better precision when looking at the detection of stimuli. The limitation of this approach, however, is that as far as most people tend to feel their heartbeats during cardiac systole, specifically approximately 200-400ms following the R-peak, there is a variability regarding when exactly within the cardiac cycle people feel their heart beating (Brener, Liu, & Ring, 1993; Clemens, 1984; Katkin, Cestaro, & Weitkunat, 1991; Wiens & Palmer, 2001; Yates, Jones, Marie, & Hogben, 1985).

For the purposes of this analysis, at subject-level, we modelled the onsets of the heartbeats at 300ms following the R peak (with duration 0ms) and the onsets of somatosensory stimulation (also with duration of 0ms). When no stimulus was present in a given epoch, the whole duration of the epoch (750ms) was modelled. The events were labelled as Hits or Misses, False Alarms or Correct Rejections, as per signal detection theory. The remaining events (confidence ratings window, button presses, response windows) were also modelled as regressors of no interest as described in the main manuscript. Additionally, we modelled the whole duration when colourful crosses were presented on the screen (2.25s) when participants were supposed to attend to their heart beating/tactile stimuli applied to their hand. Black fixation cross presented during inter-trial interval served as an implicit baseline. All pre-processing and analyses steps followed exactly the same steps as detailed in the main manuscript.

There were three contrasts of interest: (1) the main effect of stimulus type (Heartbeats vs Somatosensory stimuli), (2) the main effect of correct signal detection (Hits vs Misses), and (3) the interaction effect (Cardiac Hits – Cardiac Misses vs Somatosensory Hits – Somatosensory Misses).

We also performed the gPPI analysis, using the same right insula seed, testing a contrast between the two interaction regressor coefficients (i.e., Cardiac Hits vs Misses x Insula ROI – Somatosensory Hits vs Misses x Insula ROI).

### Supplementary results

We found a main effect of stimulus type, with greater activation in the right lateral occipital cortex while processing heartbeats compared to somatosensory stimuli (SFigure 3, STable 2). There was also the main effect of detection: Hits compared to Misses evoked greater activations encompassing cortical (frontal, parietal and occipital) as well as subcortical areas bilaterally across both focus conditions (SFigure 4, STable 3). These included precentral gyri, inferior, middle and superior frontal gyri, paracingulate cortex, insula, thalamus, putamen and caudate, brain stem, supramarginal gyrus, superior parietal lobule, postcentral gyri, lateral occipital cortex and precuneus. The reversed contrast (Misses > Hits), revealed an activation encompassing cortical (frontal, temporal and occipital) as well as subcortical areas bilaterally (SFigure 4, STable 3). These include frontal pole, cingulate and paracingulate gyrus, precuneus, intracalcarine cortex, cuneal cortex, occipital pole, lingual gyrus, parahippocampal gyrus, hippocampus, middle and inferior temporal gyrus. However, there was no stimulus type by detection interaction effect. Overall, these results are highly consistent with the findings from our main analysis.

PPI results, looking at the stimulus type by detection interaction, were also similar to the findings in the main analysis, but more confined (SFigure 5, STable 4). Right insula showed greater functional connectivity with the right lateral occipital cortex extending towards middle temporal gyrus as well as occipital pole extending to supracalcarine and intracalcarine cortex.

Therefore, overall, our findings using both approaches are highly similar and our conclusions stand.

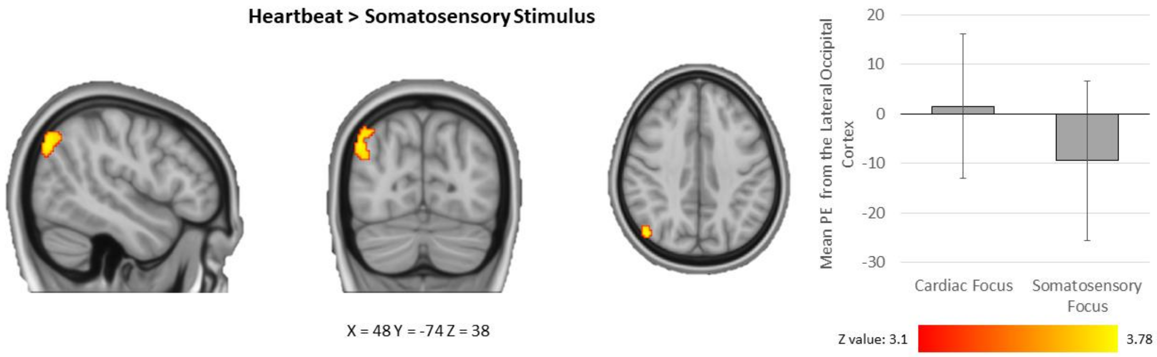


*SFigure 4 The main effect of Stimulus type showing regions presenting greater activations in the Cardiac Focus vs Somatosensory Focus condition. All images are presented in the radiological convention: the right side of the brain is depicted in the left side of the image with coordinated in the MNI space. Error bars represent standard error of the mean. PE – parameter estimate.*

*STable 2 The main effect of stimulus type: Heartbeat > Somatosensory. The coordinates for clusters maxima are presented in MNI space.*

| Cluster size (voxels) | *P* | Z-max | Coordinates | | | Side | Peak activation region |
|---|---|---|---|---|---|---|---|
| | | | X | Y | Z | | |
| 186 | .044 | 3.78 | 48 | -74 | 38 | Right | Lateral occipital cortex |

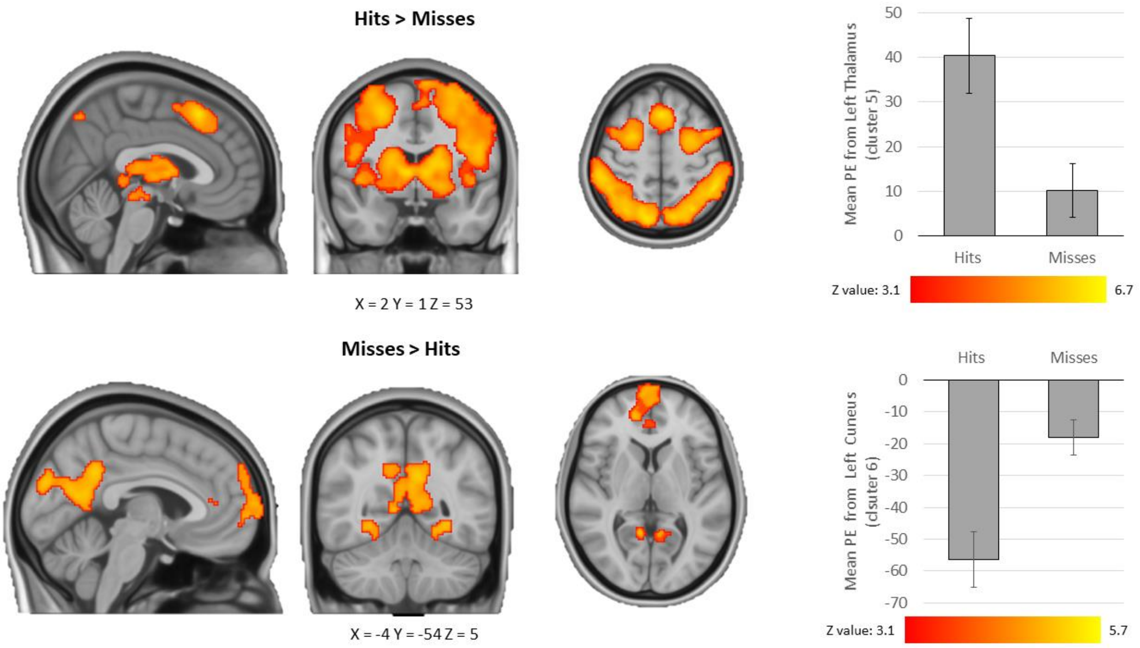


*SFigure 5 Results showing the main effect of detection. All images are presented in the radiological convention: the right side of the brain is depicted in the left side of the image with coordinated in the MNI space. Error bars represent standard error of the mean. PE – parameter estimate.*

*STable 3 Main effect of detection. The coordinates for clusters maxima are presented in MNI space.*

| Cluster size (voxels) | *P* | Z-max | Coordinates | | | Side | Peak activation region |
|---|---|---|---|---|---|---|---|
| | | | X | Y | Z | | |
| **Hits > Misses** | | | | | | | |
| 24182 | < .001 | 6.34 | -6 | 22 | 44 | Left | Thalamus |
| 11322 | < .001 | 6.64 | -28 | -68 | 56 | Left | Lateral occipital cortex |
| 594 | .001 | 5.54 | 30 | -66 | -24 | Right | Cerebellum |
| 395 | .005 | 5.00 | 54 | -34 | -14 | Right | Inferior temporal gyrus |
| 392 | .005 | 5.39 | -26 | -70 | -22 | Left | Cerebellum |
| **Misses > Hits** | | | | | | | |
| 2784 | < .001 | 5.51 | 12 | -86 | 30 | Right | Cuneal cortex |
| 1310 | < .001 | 4.78 | -8 | 58 | 36 | Left | Frontal pole |
| 740 | < .001 | 4.89 | -28 | -44 | -14 | Left | Temporal fusiform cortex |
| 651 | < .001 | 4.51 | 28 | -50 | -8 | Right | Temporal fusiform cortex |
| 528 | .001 | 5.68 | -50 | 0 | -24 | Left | Middle temporal gyrus |
| 296 | .018 | 4.45 | 40 | 6 | -30 | Right | Temporal pole |

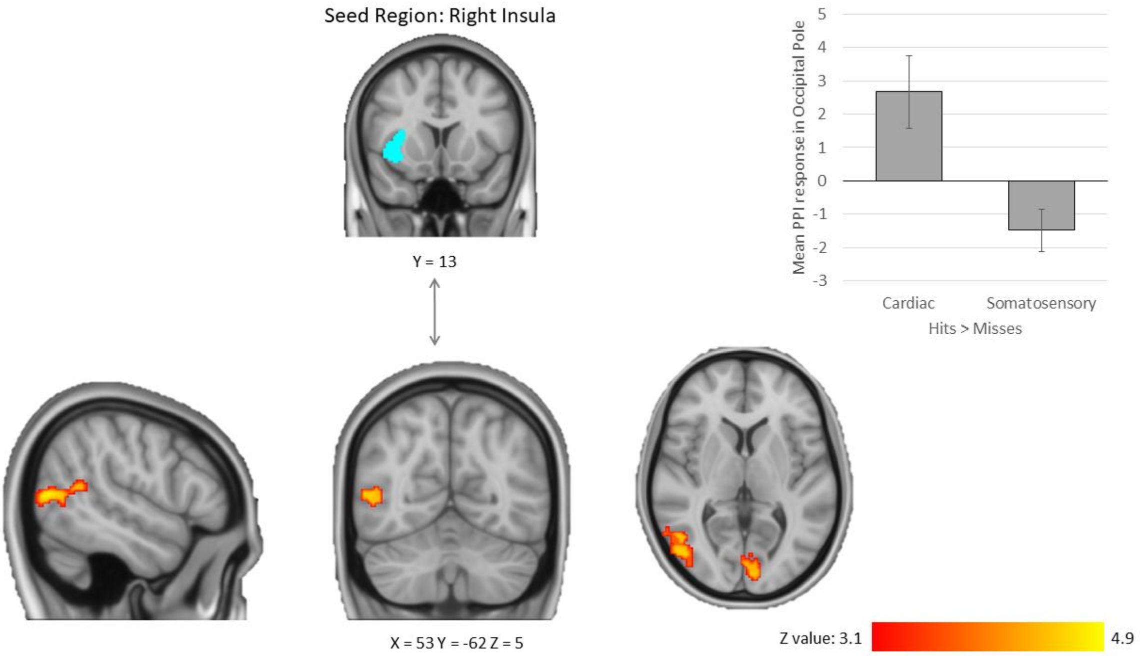


*SFigure 6 PPI Analysis showing greater functional connectivity between the right insula seed and occipital areas in the Correct detections (Hits > Misses) during Cardiac vs Somatosensory condition. All images are presented in the radiological convention: the right side of the brain is depicted in the left side of the image with coordinated in the MNI space. Error bars represent standard error of the mean.*

*STable 4 Results of the PPI Analysis. The coordinates for clusters maxima are presented in MNI space.*

| Cluster size (voxels) | *P* | Z-max | Coordinates | | | Side | Peak activation region |
|---|---|---|---|---|---|---|---|
| | | | X | Y | Z | | |
| 284 | .005 | 4.85 | 54 | -70 | 8 | Right | Lateral occipital cortex |
| 264 | .008 | 3.95 | -6 | -90 | 6 | Left | Occipital pole |
